# The logical structure of experiments lays the foundation for a theory of reproducibility

**DOI:** 10.1101/2022.08.10.503444

**Authors:** Erkan O. Buzbas, Berna Devezer, Bert Baumgaertner

**Author notes:** These authors contributed equally to this work.

## Abstract

The scientific reform movement has proposed openness as a potential remedy to the putative reproducibility or replication crisis. However, the conceptual relationship between openness, replication experiments, and results reproducibility has been obscure. We analyze the logical structure of experiments, define the mathematical notion of idealized experiment, and use this notion to advance a theory of reproducibility. Idealized experiments clearly delineate the concepts of replication and results reproducibility, and capture key differences with precision, allowing us to study the relationship among them. We show how results reproducibility varies as a function of: the elements of an idealized experiment, the true data generating mechanism, and the closeness of the replication experiment to an original experiment. We clarify how openness of experiments is related to designing informative replication experiments and to obtaining reproducible results. With formal backing and evidence, we argue that the current “crisis” reflects inadequate attention to a theoretical understanding of results reproducibility.

## 1 Introduction

In a number of scientific fields, replication and reproducibility *crisis* labels have been used to refer to instances where many results have failed to be corroborated by a sequence of scientific experiments. This state of affairs has led to a scientific reform movement. However, this labeling is ambiguous between a crisis of practice and a crisis of conceptual understanding. Insufficient attention has been given to the latter, which we believe is a detriment to moving forward to conduct science better. In this paper, we make theoretical progress toward understanding replications and reproducibility of results (henceforth “results reproducibility”) by a formal examination of the logical structure of experiments^1^.

We view replication and reproducibility as methodological subjects of metascience. As we have emphasized elsewhere (Devezer et al., 2021), these methodological subjects need a formal approach to properly study them. Therefore, our work here is necessarily mathematical; however, we make our conclusions relatable to the broader scientific community by pursuing a narrative form in explaining our framework and results within the main text. Mathematical arguments are presented in the appendices. Our objective is to build a strong, internally consistent, verifiable theoretical foundation to understand and to develop a precise language to talk about results reproducibility. We advance mathematical arguments from first principles and proofs, using probability theory, mathematical statistics, statistical thought experiments, and computer simulations. We ask the reader to evaluate our work within its intended scope of providing theoretical precision and nuanced arguments.

The following backdrop to motivate our research matters: A common concern voiced in the scientific reform literature and recent scholarly discourse regards various forms of scientific malpractice as potential culprits of reproducibility failures and openness is sometimes touted as a remedy to alleviate such malpractices (Collins and Tabak, 2014; Iqbal et al., 2016; National Academies of Sciences, Engineering, and Medicine, 2017; Nosek et al., 2015, 2022). Some malpractice is believed to take place at the level of the scientist. For example, hypothesizing after the results are known involves presenting a post hoc hypothesis as if it were an a priori hypothesis, conditional on observing the data (Kerr, 1998; Munafò et al., 2017). Another example is p-hacking, a statistically invalid form of performing inference to find statistically significant results (Bruns and Ioannidis, 2016; Gelman and Loken, 2013; Munafò et al., 2017). Some is believed to operate at the community or institution level. For example, publication bias involves omitting studies with statistically nonsignificant results from publications and is primarily attributed to flawed incentive structures in scientific publishing (Collaboration et al., 2015; Munafò et al., 2017). Before we suspect malpractice of either kind and set out to correct the scientific record or demand reparations, however, it behooves the scientific community to gain a complete understanding of the factors that may account for a given sequence of research results.

If a result of an experiment is not reproduced by a replication experiment, before we reject it as a false positive or suspect some form of malpractice, we need to assess and account for: i) sampling error, ii) theoretical constraints on the reproducibility rate of the result of interest, conditional on the elements of the original experiment, and iii) assumptions from the original experiment that were not carried over to the replication experiment. First of these is a well-known and widely understood statistical fact that describes why methodologically we can at best guarantee reproducibility of a result on average (that is, in expectation). The second point about the theoretical limits of the reproducibility rate is not well understood and we hope to address this oversight in this paper. The last one has been brought up in individual cases but typically in an ad hoc manner and we aim to provide a systematic approach for comprehensive evaluations of replication experiments. Since metascientific heuristics may lead us astray in these assessments, we need a fine-grained conceptual understanding of how experiments operate and relate to each other, and what role openness plays in facilitating replications or promoting reproducible results. Indeed a replication crisis and a reproducibility crisis are different things, and should be understood on their own. We distinguish between replication experiments and results reproducibility, and discuss precursors of each.

In this paper, we argue that “failed” replications do not necessarily signify failures of scientific practice^2^. Rather, they are expected to occur at varying rates due to the features of and differences in the elements of the logical structure of experiments. Using a mathematical characterization of this structure, we provide precise definitions of and clear delineation between replication, reproducibility, and openness. Then, using toy examples, simulations, and cases from the scientific literature, we illustrate how our characterization of experiments can help identify what makes for replication experiments that can, in theory, reproduce a given result and what determines the extent to which experimental results are reproducible. In the next section, we define main notions that we use to build a logical structure of experiments which helps us derive our theoretical results.

## 2 The logical structure of experiments

### 2.1 Definitions

The *idealized experiment* is a probability experiment: A trial with uncertain outcome on a well-defined set. A scientific experiment where inference is desired under uncertainty can be represented as an idealized experiment. The results from an experiment can be defended as valid only if the assumptions of the probability experiment hold. One useful setup for us is as follows: Given some background knowledge *K* on a natural phenomenon, a scientific theory makes a prediction, which is in principle testable using observables, the data *D*. A mechanism generating *D* is formulated under uncertainty and is represented as a probability model *M*_*A*_ under assumptions *A*. Given *D*, inference is desired on some unknown part of *M*_*A*_. The extent to which parts of *M*_*A*_ that are relevant to the inference are confirmed by *D* is assessed by a fixed and known collection of methods *S* evaluated at *D* (similar descriptions for other purposes can be found in Devezer et al., 2019, 2021).

#### Definition 1.

The tuple *ξ* := (*K, M*_*A*_, *S, D*) is an *idealized experiment*.

Definition 1 of *ξ* captures some key distinct elements of experiments whose population characteristics can in principle be tested. These elements are not necessarily independent of each other. For example, *K* may inform and constrain the sets of plausible *M*_*A*_ and *S*. Or it may be necessary for *M*_*A*_ to constrain *S*.

*M*_*A*_ includes the sampling design when sampling a population conforming *A*, which we assume to be independent of sampling design. For example, *A* may be the description of an infinite population of interest, which may be sampled in a variety of ways to yield distinct probability models *M*_*A*_ for the data depending on the sampling scheme.

We distinguish two elements of *S*: *S*_*pre*_ and *S*_*post*_. *S*_*pre*_ is the *scientific* methodological assumptions made prior to data collection and procedures implemented to obtain *D. S*_*pre*_ captures assumptions in designing and executing an experiment such as experimental paradigms, study procedures, instruments, and manipulations.

Conditional on *K* and *M*_*A*_, *S*_*pre*_ is *reliable* if the random variability in *D* is due only to sampling variability modeled by *M*_*A*_. *S*_*post*_ is the *statistical* methods applied on *D*. If inferential, *S*_*post*_ is *reliable* if it is statistically consistent. *S* is reliable if and only if *S*_*pre*_ and *S*_*post*_ are reliable.

We also distinguish two elements of *D*: *D*_*s*_ and *D*_*v*_. *D*_*s*_ is the structural aspects of the data, such as the sample size, number of variables, units of measurement for each variable, and metadata. *D*_*v*_ is the observed values, that is, a realization conforming *D*_*s*_. Some statistical approaches to assess risk and loss focus on the reproducibility conditional on *D*_*v*_, whereas others focus on averages over independent realizations of *D*_*v*_.

Definition 1 of *ξ* allows us to scaffold other definitions as follows. An exact *replication experiment ξ′* must generate *D′* independent of *D* conditional on *M*_*A*_ in the values but with the same structure *D*_*s*_.

#### Definition 2.

The tuple 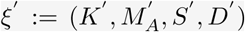 is an *exact replication experiment* of *ξ* if 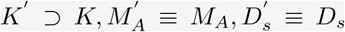 and 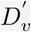 is a random sample independent of *D*_*v*_. If at least one of 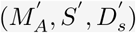 differs from (*M*_*A*_, *S, D*_*s*_) or *K′* ⊅ *K*, then *ξ′* can at most be a *non-exact replication experiment* of *ξ*.

Definition 2 mathematically isolates *ξ* and *ξ′* from *R*. That is, *ξ′* does not need to have a specific aim to be performed or worked with as a mathematical object. The benefits of this isolation will become clear in section 3, where an unconditional *ξ* and its *non-exact ξ′* pair may become a *ξ* and its *exact ξ′* pair, conditional on *R*.

Often, however, we would perform experiments with a specific aim and would like to see whether the result of *ξ* is reproduced in *ξ′*. Depending on the desired mode of statistical inference, example aims include hypothesis testing, point or interval estimation, model selection, or prediction of an observable. Further, when augmented with an *R, K′* must differ from *K* in a specific way. Encompassing all these statistical modes of inference, we introduce the notion of a *result R*, as a decision rule. For convenience, we assume that *R* lives on a discrete space here.

#### Definition 3.

Let 𝒳 be the sample space and ℛ ≡ {*r*_1_, *r*_2_, …, *r*_*q*_}, *q* ∈ ℤ^+^ be the decision space. For sample size *n* ∈ ℤ^+^, the function *R* : 𝒳 ^*n*^ → ℛ is a *result*.

*R* is obtained by mapping the application of *S*_*post*_ on *D* on to the decision space. If *ξ′* is aimed at reproducing *R* of *ξ*, it is conditional on *R* and leads us to the following connection between an idealized experiment and a result.

#### Definition 4.

Let *R* and *R′* be results from *ξ* and *ξ′* respectively. *R* = *r*_*o*_ is *reproduced* by *R′* = *r*_*d*_ if *d* = *o*, else *R* = *r*_*o*_ is *not reproduced*.

In definition 4, reproducibility of *R* depends on the available actions *r*_1_, *r*_2_,, *r*_*q*_. The size of *q* is case specific. Examples are as follows. In a null hypothesis significance test, *q* = 2 : the null hypothesis and the alternative hypothesis. In a model selection problem we entertain *q* models and choose one as the best model generating the data. In a parameter estimation problem for a continuous parameter, we build *q* arbitrary bins, and call a result reproduced if the estimate from *ξ′* falls in the same bin as the result from *ξ*. How the bins are constructed in a problem affects the actual reproducibility rate of a result. However, for our purposes in this paper, theoretical results hold for all cases regardless of this tangential issue.

The class of problems of interest to us here involves cases where in a *sequence* of exact replication experiments, if *S* is reliable, we should expect a regularity in the results. That is, probability theory tells us that if the elements of an idealized experiment are well-defined, then we should expect the results from a sequence of replication experiments to stabilize at a certain proportion, given the characteristics of an idealized experiment and the true data generating mechanism. This notion is formalized in definition 5.

#### Definition 5.

Let *ξ*^(1)^, *ξ*^(2)^, …, *ξ*^(*N*)^ be a sequence of idealized experiments. The *reproducibility rate*

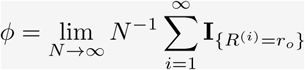

of a result *R* = *r*_*o*_ is a parameter of the sequence (**I**_{*C*}_ = 1 if *C*, and 0 otherwise).

An advantage of definition 5 is that conditional on *R* = *r*_*o*_ in *ξ* and a sequence of replication experiments *ξ*^(1)^, *ξ*^(2)^, …, *ξ*^(*N*)^, the *relative frequency* of reproduced results *ϕ*_*N*_ converges to *ϕ* ∈ [0, 1] as *N* → ∞. So, we immediately have 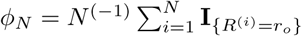 as a natural estimator of *ϕ*. Further, we are formally comforted to know that lim_*N*→∞_ ℙ (*ϕ*_*N*_ = *ϕ*) = 1. That is, with high probability, the estimated reproducibility rate *ϕ*_*N*_ from a sequence of replication experiments will get closer to the true reproducibility rate of the original experiment *ϕ*.

Finally, we turn to the last of our key concepts: *openness*. Openness refers to the accessibility of all necessary information regarding the elements of *ξ* by another idealized experiment *ξ*^*^. This accessibility may be used for a variety of purposes. For example, *S*_*post*_ can be re-applied to *D* to verify *R* independently of *ξ*. In this capacity, openness facilitates the auditing of experimental results by way of screening off certain errors, including human and instrumental (e.g., data entry and programming errors), that may be introduced in the process of obtaining *R* initially. On the other hand, openness may be needed to perform an exact *ξ′* by way of duplicating *S*_*pre*_ to obtain *D′* and *S*_*post*_ to obtain *R′*. In this capacity, openness makes exact *ξ′* possible.

Openness is critically related to reproducibility since the degree to which information is transferred from *ξ* to *ξ′* impacts the *ϕ* of a given result. However, not all elements of *ξ* need to be open for all purposes. Therefore, a nuanced understanding of openness requires evaluating it at a fixed configuration of the elements of *ξ* conditional on a specific purpose, rather than as a categorical judgment at the level of the whole experiment, as open or not. This leads us to think of openness element-wise, as in definition 6.

#### Definition 6.

Let Π be the power set of elements of *ξ* and *π* ∈ Π. *ξ* is *π*-*Open* for *ξ*^*^ if *π* ⊂ *K*^*^ where *ξ*^*^ is an idealized experiment that imports information from *ξ*.

A specific example of *π*-*Open* of definition 6 would be *π* ≡ (*M*_*A*_, *S*_*pre*_) where *ξ*^*^ gets all the information about the assumptions, model, pre-data methods from *ξ* but no other information. Another example of *π*-*Open* is the special case where *ξ* has all its elements open, that is *π* ≡ (*K, M*_*A*_, *S, D*). In this case, for convenience, we say *ξ* is *ξ*-*Open* for *ξ*^*^.

### 2.2 Fundamental results on replications and reproducibility rate from first principles

Here we present two results about reproducibility and some remarks, based on definitions 1-6. A well-formed theory of reproducibility requires results of these type: fundamental, mathematical, and invoking a functional framework to study replications and reproducibility. They serve as theoretical benchmarks to check other results against. Technically oriented readers may refer to Appendix 1 and Appendix 2 for a more detailed discussion and results complementary to the main argument.

We begin by noting that, given definition 5 and the discussion following it, it is not straightforward to say exactly what we gain if we were to update the estimated reproducibility rate based on the results obtained from performing more replications. Indeed, to understand the value of replication experiments in assessing the reproducibility of a result, a strong mathematical statement is required, which is our result 1.

#### Result 1.

Let *ξ*^(1)^, *ξ*^(2)^, …, *ξ*^(*N*)^ be a sequence of replication experiments with reproducibility rate *ϕ* given by definition 5. Then,

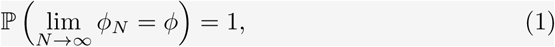

where *ϕ*_*N*_ is the sample reproducibility rate of result *R* = *r*_*o*_ obtained from the sequence (proof in Appendix 1).

Result 1 is fundamental to study replications and reproducibility for a number of reasons:

1. It provides a basis for building trust in the notion of reproducibility from replication experiments. Roughly, it says that if we perform replication experiments and estimate the reproducibility rate of *r*_*o*_ by *ϕ*_*N*_ from these experiments, then we are *guaranteed* that deviations of *ϕ*_*N*_ from *ϕ* are going to *get small* and *stay small*.
2. It is almost necessary to move forward theoretically. It immediately implies that if the assumptions of an original experiment are satisfied in its replication experiments, then we are *adopting a statistically defensible strategy* by continuing to perform replication experiments and updating *ϕ*_*N*_ as a proportion of successes to assess the reproducibility rate. Therefore, result 1 gives us a theoretical justification of *why we should care* about performing more replication experiments whose assumptions are satisfied and be interested in estimating reproducibility rate based on those replication experiments alone. Further, violating the assumptions of *ξ* in replication experiments implies that *ϕ*_*N*_ converges to some *ϕ* defined by the flaws underlying a non-exact sequence of replications of *ξ* rather than the reproducibility rate of *r*_*o*_ of interest.
3. As we will detail in result 2, a theoretically fertile way to study replication experiments is by defining a sequence of experiments as a stochastic process. The results from such processes almost always require the solid foundation provided by result 1.

#### Remark 1.

The reproducibility rate given in definition 5 has excellent properties as shown by result 1. However, we keep in mind that definition 5 is only one way to measure reproducibility. It is a counting measure which counts the reproduced results. Instead, a continuous measure as a degree of confirmation of a result might seem more proper to measure reproducibility. One has to be aware that just defining a reproducibility measure does not imply that it has desirable mathematical properties. It is easy to define meaningful continuous measures of reproducibility which might have pathological properties (e.g., that do not satisfy result 1) and these should be avoided (see Appendix 1 for details).

In practice *S*_*post*_ are functions of sample moments, such as the sample mean. In these cases, sometimes the Lindeberg-Lévy Central Limit Theorem (CLT) and its extensions provide useful results about the properties of *ξ*^(1)^, *ξ*^(2)^, …. However, restricting *S*_*post*_ this way constrains the mathematical setting to study the statistical properties of *ξ*^(1)^, *ξ*^(2)^, … or results reproducibility. For example, working with the CLT is challenging when *S*_*post*_ cannot be formulated as a function of a fixed sample size or to discuss the properties of a sequence of replication experiments directly, without referring to *S*_*post*_ as a means to estimate a particular *R*.

We provide a broad setting without these limitations by assuming that *K* requires only minimal validity conditions on *M*_*A*_ and *S*. Specifically, we let *M*_*A*_ be any probability model, subject only to some mathematical regularity conditions such as continuity of distribution functions, the existence of the mean and the variance of the variable of interest. We also let *S*_*post*_ be the sample distribution function^3^. With the generality provided by these assumptions, we obtain one of our main theoretical results.

#### Result 2.

The sequence of idealized experiments *ξ*^(1)^, *ξ*^(2)^, … given by definition 5 is a proper stochastic process, seen as a joint function of random sample *D* and of each value in the support of data generating mechanism, *x* ∈ ℝ (see constructive proof in Appendix 2).

Result 2 is of fundamental importance to study results reproducibility mathematically because it allows us to apply the well-developed theory of stochastic processes to build a theory of results reproducibility. Two aspects of result 2 are noteworthy:

1. When we obtain a random sample in *ξ* and perform inference using a fixed value of a statistic such as a threshold, the sequence *ξ*^(1)^, *ξ*^(2)^, … constitutes random variables independent of each other conditional on the true model generating the data. Obtaining the distributions implied by *ξ* helps us understand the statistical nature of replication experiments.
2. *ξ′* generates new data *D′* and *R′* is conditional on *D′*. That is, when inference is performed for a particular replication experiment, the data are fixed. Most generally, conditional on *D′* if the empirical distribution function is *R′*, then the replication experiment estimates the model generating the data. Therefore, a replication experiment determines a sample based estimate of a statistical model.

Summary of all notation and terms introduced in this section can be found in table 1 for quick reference. In the next section, we introduce a toy example as a running case study to instantiate our theoretical results on replications, reproducibility, and openness.

**Table 1.**
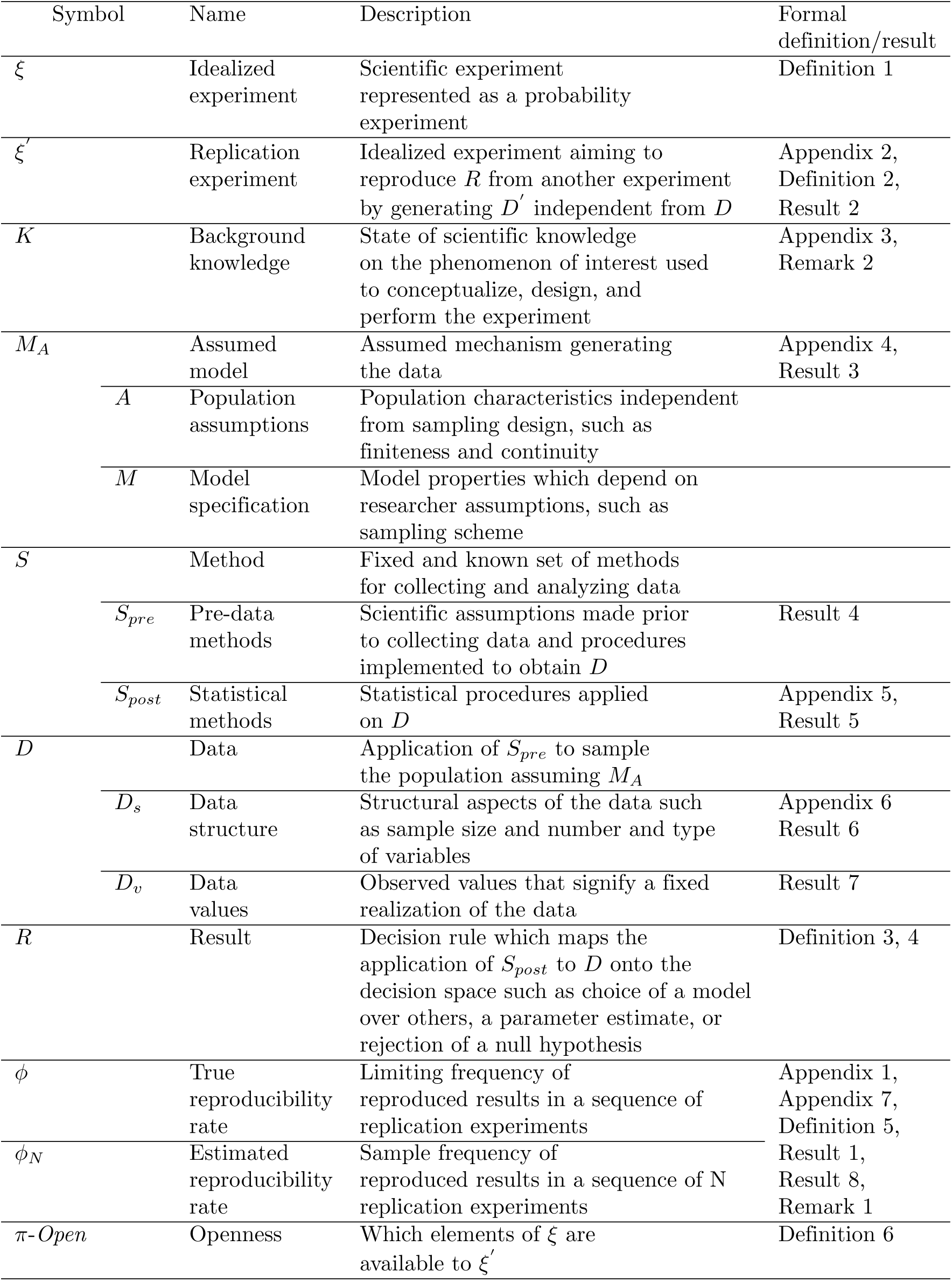
Quick reference guide to notation and key terms.

## 3 A toy example

Our toy example involves an inference problem regarding a population of ravens, *K*. An infinite population of ravens where each raven is either black or white constitutes the population assumptions, *A*. Each uniformly randomly sampled raven can be identified correctly as black or white, which defines the pre-data methods, *S*_*pre*_. The result of interest, *R*, is to estimate the (unknown) population proportion of black ravens, *p*, or some function of it.

We consider six distinct sampling scenarios, which lead to six distinct *M*_*A*_, and thus six distinct idealized experiments. To avoid overly complicated mathematical notation we denote the models by: *ξ*_*bin*_, *ξ*_*negbin*_, *ξ*_*hyper*_, *ξ*_*poi*_, *ξ*_*exp*_, *ξ*_*nor*_. These models represent the binomial, negative binomial, hypergeometric, Poisson, exponential, and normal probability distributions for the data generating mechanism, respectively. In specific examples, we also vary *S*_*post*_, the point estimator of the parameter of interest to take values as maximum likelihood estimate (MLE), method of moments estimate (MME), and posterior mode (i.e., Bayesian inference). We further vary *D*_*s*_ via the sample size (i.e., *n* ∈ {10, 30, 100, 200}). We use these idealized experiments to illustrate our results in the rest of the paper.

These six idealized experiments make the following sampling assumptions. *ξ*_*bin*_ stops when *n* ravens are sampled. *ξ*_*negbin*_ stops when *w* white ravens are sampled. *ξ*_*hyper*_ is a special case where the sampling has access only to a finite subset of the infinite population delineated by *A. ξ*_*bin*_, *ξ*_*negbin*_, *ξ*_*hyper*_ are often called *exact* models, in the sense that their *M*_*A*_ does not involve any limiting or approximating assumptions. On the other hand, *ξ*_*poi*_ approximates *ξ*_*bin*_ where a large sample of *n* ravens is sampled when the proportion of black ravens *p* is small. The larger the *n* and the smaller the *p* such that *np* remains constant, the better the approximation. *ξ*_*exp*_ has the same approximative characteristics and parameter as *ξ*_*poi*_. However, notably, *ξ*_*exp*_ records the time between observations instead of counting the ravens, so its *S*_*pre*_ is different from all other experiments. Finally, *ξ*_*nor*_ approximates *ξ*_*bin*_ where a large sample of *n* ravens with intermediate proportion of black ravens, *p*, holds.

As the result of interest, *R*, these six idealized experiments aim to estimate either the proportion of black ravens, *p*, in the population or the rate of black ravens sampled, *np* → *λ*, a function of *p*, in the approximative models. Figure 1 shows distinctive elements of these six idealized experiments.

**Figure 1.**
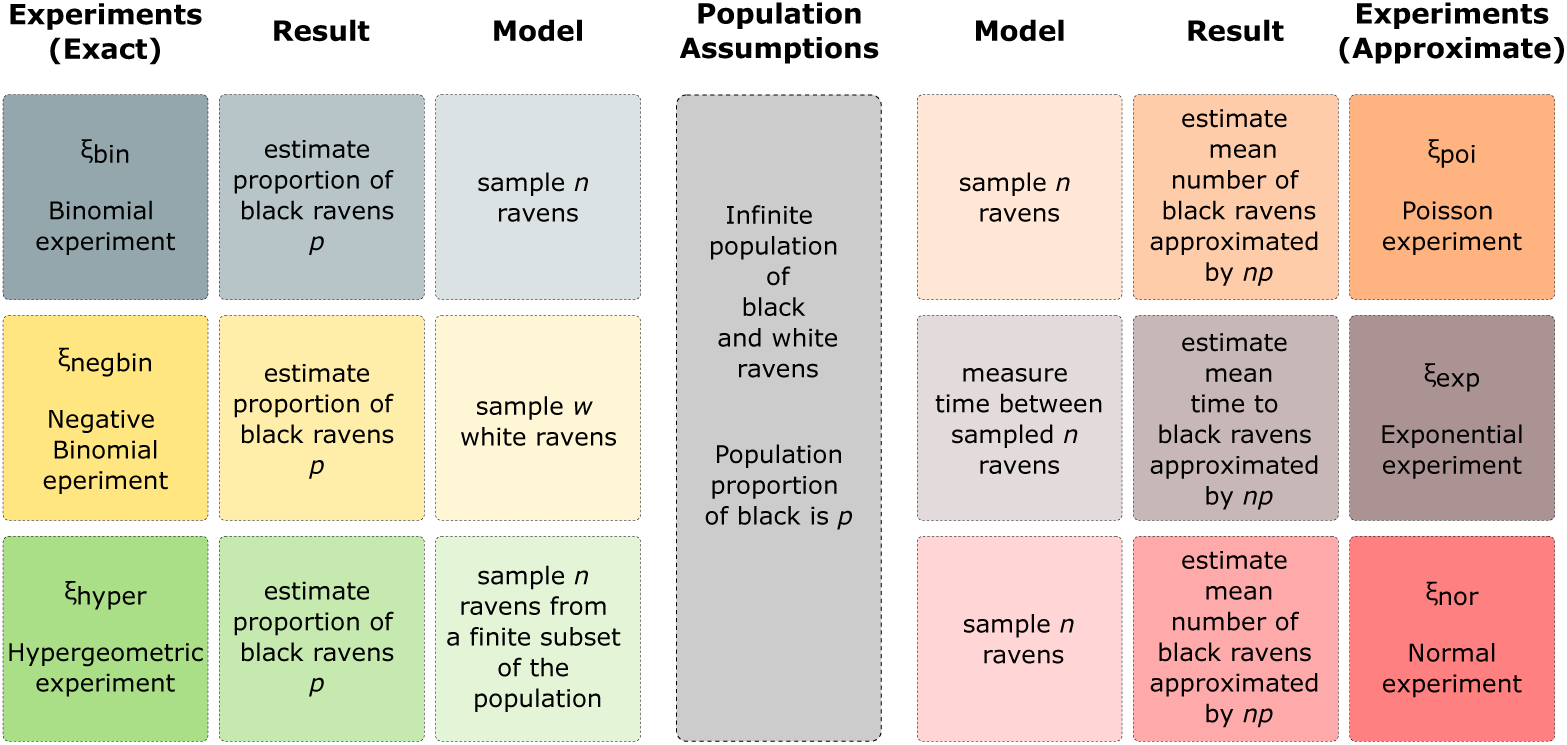
Six idealized experiments *ξ*_*bin*_, *ξ*_*negbin*_, *ξ*_*hyper*_, *ξ*_*poi*_, *ξ*_*exp*_, *ξ*_*nor*_ : The binomial, negative binomial, hypergeometric, Poisson approximation to binomial, exponential waiting times between Poisson events, and normal approximation to binomial, respectively. All but *ξ*_*hyper*_ assume infinite population (*A*) of black and white ravens, with sampling designs resulting in distinct probability models (*M*_*A*_). *ξ*_*hyper*_ assumes sampling from a finite subset of the population. All experiments aim at performing inference on result (*R*), which reduces down to an estimate of either the population proportion of black ravens or the mean number of black ravens in the population.

In section 4, we use these six idealized experiments to show that *openness* connects to reproducibility in a variety of ways and to *reproduce* a given result, *replication experiments* do not need to be *exact*. We show that conditional on a given result from an original experiment, *non-exact* replication experiments can serve as valid *exact* replication experiments, if the inferential equivalence holds between the original and the replication. We further show that, the true rate of reproducibility of a sequence of exact replication experiments and a sequence of non-exact replication experiments are distinct (except trivially) for a given result.

## 4 Element-wise openness and assessing the meaning of replications

Tools and procedures have been developed to help facilitate openness in science (Collins and Tabak, 2014; Munafò et al., 2017; Nosek et al., 2015; Wagenmakers et al., 2012). Guidelines may argue for making as much information available as possible about an experiment or leave it to intuition to guide which elements of an experiment are relevant and need to be shared for replication. We are interested in better understanding what does and does not need to be made available, in service of which objective, and under what conditions. We perceive two main issues: what openness means for performing meaningful replications and how it impacts results reproducibility. We first evaluate the former. Then we show that a uniform, wholesale framing of openness is not the remedy to the reproducibility crisis that some take it to be.

*ξ* has elements involving uncertainty, such as *D*_*v*_ taken as a random variable. Uncertainty modeled by probability is always conditional on the available background information (Lindley, 2000), and thus reproducibility of *R* is always conditional on *K*. That is, *ξ′* must import sufficient information from *ξ* with respect to *R* of interest to assess whether *R* is reproduced in *R′*. A *ξ′* that aims to reproduce a given result from *ξ* may be performed in a variety of ways depending on which elements of *ξ* are open.

In the context of our toy example, figure 2 shows a network structure of some possible *ξ* as a function of which elements of *ξ* are open. Specifically, we consider variations of the six experiments introduced in section 3 for two *S*_*post*_ (MLE and posterior mode) and two *D*_*s*_ (*n* = 30 and *n* = 200) yielding 24 distinct *ξ*, each denoted by a node in each network in figure 2. Given one of these 24 as *ξ*, all possible 24 experiments are either exact or non-exact *ξ′*. We use definition 6 and specify *π* to assess the degree of openness in these experiments. When *ξ* is *ξ*-*Open*, the probability of exact replication is 1 and every node of the network is only connected to itself. If *ξ* is *π*-*Open*, where *π* is a proper subset of *ξ* then *ξ′* may be a non-exact replication of *ξ* in various ways because *ξ′* needs to substitute in a value for elements that are not in *π*. Therefore, the probability of *ξ′* being an exact replication of *ξ* is lower than when *ξ* is *ξ*-*Open*. In figure 2, we show the network structures that result from choosing non-open elements with equal probability among all substitutions considered for each element. The network complexity depends on the size of *π*. If it is large, the number of connections among the nodes in the network is small and each connection is strong (e.g., strongest when all open). In contrast, if it is small, the number of connections among the nodes in the network is large because there are both multiple substitutions to be made and multiple possibilities for each, and each connection is weak (e.g., weakest when *M*_*A*_, *S*_*post*_, *D*_*s*_ not open in figure 2). Hence, as the size of *π* decreases, it becomes less probable to perform an exact replication of *ξ*. By looking at which elements of *ξ* are open to start with, we can assess how the sequence *ξ*^(1)^, *ξ*^(2)^, … of replication experiments can be misinterpreted if the necessary elements were not open and/or got lost in translation. In the rest of this section, we organize our results by elements *K, M*_*A*_, *S, D*.

**Figure 2.**
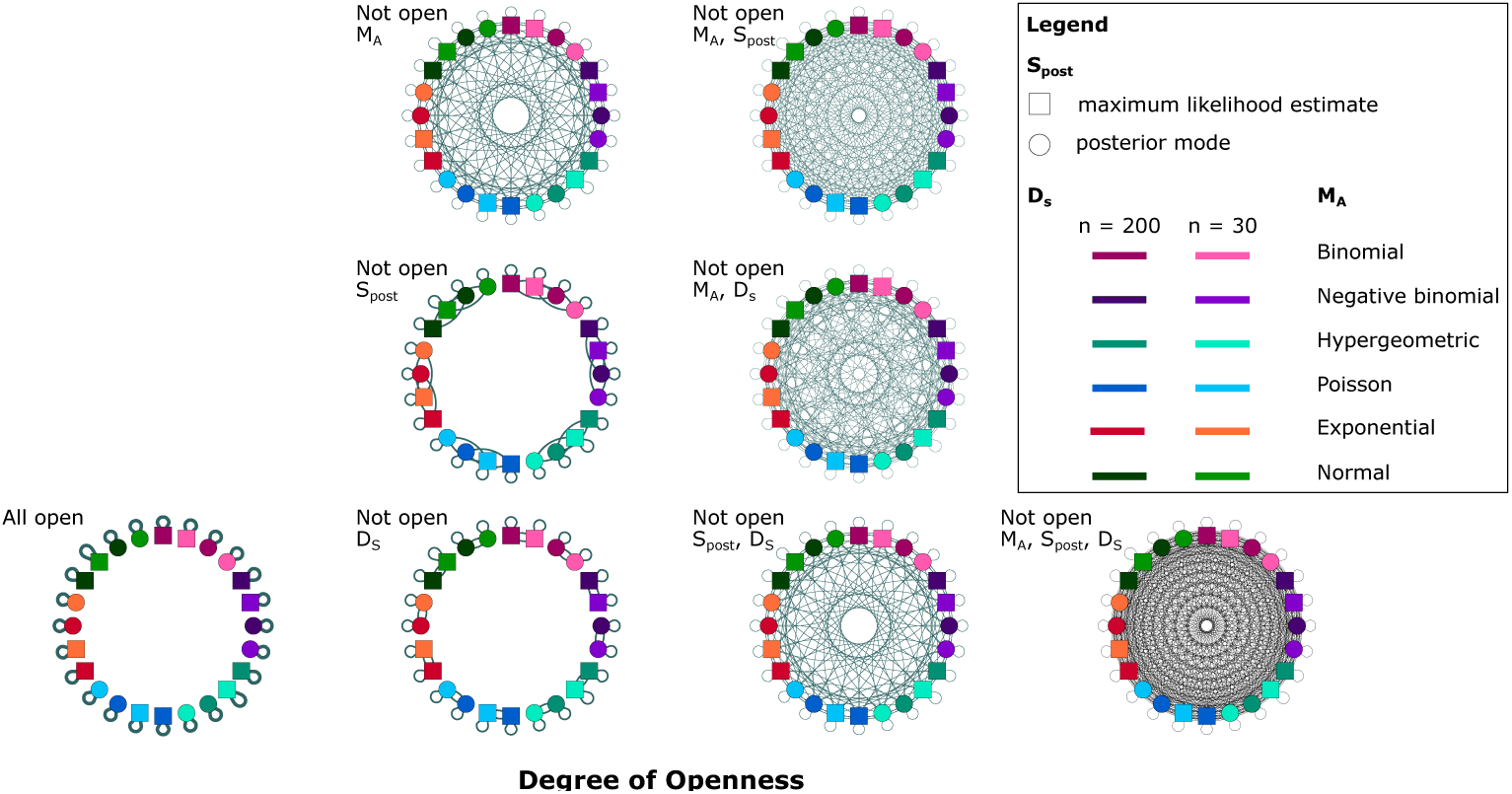
For the models in the toy example, degrees of openness (as given by definition 6) are depicted in 8 networks, each consisting of the same 24 idealized experiments. Each idealized experiment is represented by a node in each network. These 24 experiments are obtained by a 6 × 2 × 2 factorial design. The first factor, *M*_*A*_, takes 6 values: binomial, negative binomial, hypergeometric, Poisson, exponential, normal. The second factor, *S*_*post*_, takes 2 values: MLE and Posterior mode. The third factor, *D*_*s*_, takes two values: *n* = 30 and *n* = 200. Connections between nodes represent potential substitutions of non-open elements of idealized experiments. As more elements of an idealized experiment are non-open, the probability of choosing an exact replication decreases, as indicated by increased connectivity in the network.

### 4.1 Background knowledge, *K*

Providing an exact description of what goes into *K* is notoriously difficult. *K*, which is more of a philosophical element of *ξ*, typically carries over much more than what can be immediately gleaned over by a transparent and complete description of *M*_*A*_, *S*, and *D*. We understand *K* to contain theoretical assumptions, contextual knowledge, paradigmatic principles, a specific language, presuppositions inherent in a given field; in short, a lot of inherited cultural and historical meaning of the kind Feyerabend refers to as *natural interpretations* in Against Method (Feyerabend, 1993, p. 49). As Feyerabend explains, such natural interpretations are not easy to make explicit or even sometimes be aware of and thus, being open about them might not be a matter of choice. However, observations gain meaning only against this backdrop and experiments can only be interpreted correctly by using the same language used to design them in the first place. Within *ξ*, this tends to happen implicitly whereas when performing *ξ′*, there is no guarantee that all the relevant information in *K* will carry over to *K′*.

Using the binomial experiment in our toy example, we can illustrate why *K* is an integral part of *ξ* and what role it plays for *ξ′*. In *ξ*_*bin*_, our aim (*R*), is to estimate the proportion of black ravens (*p*) in an infinite population of ravens (*A*). *M*_*A*_ samples *n* ravens. As our *S*_*pre*_, we count black and white ravens by naked eye. As our *S*_*post*_, we use the maximum likelihood estimator of *p*. We set *n* = 100, which constitutes our *D*_*s*_. This description of *ξ*_*bin*_ based on a specific configuration of *M*_*A*_, *S*_*pre*_, *S*_*post*_, *D*_*s*_ could just as well be used to define an experiment in which scientists are interested in estimating the proportion of black *swans* in a population of black and white swans. While *ξ*_*bin*_ would still be mathematically well-defined, its scientific content and context is not captured by any of these four elements. For that, we need *K*. Without *K*, we would have to consider an 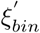 about black swans as an acceptable replication of *ξ*_*bin*_ about black ravens, based on mathematical structure alone. *K*, then, communicates scientific meaning across experiments.

As a more practical example of the import of *K*, we consider a recent “failed” replication experiment. Murre (2021) attempted to replicate a classical experiment by (Godden and Baddeley, 1975) on context-dependent memory. Context-dependent memory refers to the hypothesis that the higher the match between the context in which a memory is being retrieved and the context in which the memory was originally encoded, the more successful the recall is expected to be. In the abstract, Murre (2021) summarizes the results of the replication experiment as follows: “Contrary to the original experiment, we did not find that recall in the same context where the words had been learned was better than recall in the other context.” Does this suggest that the results of the original experiment were a false positive—as replication failures are commonly interpreted? There are many reasons to not jump at that conclusion including sampling error and the fact that the context of the replication was different from that of the original Godden and Baddeley (1975) experiment. Specifically, unlike the original, the replication was being filmed as part of a TV program. We will set these obvious concerns aside for a moment to focus on another. Ira Hyman explains the issue in a Twitter thread (Hyman, 2021). Hyman indicates that the phenomenon of context-dependent memory is conditional on the distinctiveness of the encoding context. That is, if distinct contexts are used over multiple trials, the chances that the context will be remembered with the encoded information increases. When the context is not distinctive enough or remains constant over trials, the effect disappears. Another known boundary condition for the phenomenon is the outcome variable: Past research has shown that this works for retrieval tasks (e.g., free recall) and not recognition. The Murre (2021) replication did not carry over these contextual details and changed the design in a way to not instigate context-dependent memory. As a result, the differences between *R* and *R′* become impossible to attribute to a single cause and fail to provide evidence that can confirm or refute the results of the original Godden and Baddeley (1975) experiment. It is even questionable whether the Murre (2021) experiment provided an appropriate test of the result of interest in the first place to be considered a meaningful or relevant replication.

This replication example on context-dependent memory appears to imply that a *ξ′* is meaningful or relevant with respect to a specific result *R*. By definition 2 and its interpretation, however, we know that mathematically it is more convenient to separate the definition of *ξ′* from *R*. It follows that there are at least two aspects of assessing the meaning and relevance of a replication.

First, while an operational definition of *K* is elusive, a useful way to think about *K* is “all the information in *ξ* that is not already in *M*_*A*_, *S*, and *D*”. At the minimum, for *ξ′* to be considered a *meaningful* replication of *ξ, K′* must import some information in *K* regarding the immediate scientific context of *ξ*. For this to hold, there is no need to invoke the notion of *R*.

Second, to assess the reproducibility of a given *R, K′* must import *relevant* information pertaining *R* from *ξ*. That is, replication experiments unconditional and conditional on *R* are not the same objects. To emphasize the difference between them, we distinguish between *in-principle* and *epistemic* reproducibility of an *R* in remark 2 (for further details, see appendix Appendix 3).

#### Remark 2.

Let *ξ* be an idealized experiment and *ξ′* be its exact replication. Conditional on *R* from *ξ, K′* is necessarily distinct from *K* for epistemic reproducibility of *R* by *R′*, but not necessarily distinct for in-principle reproducibility of *R* by *R′*.

In practice, *ξ′* can never be an *exact* replication of *ξ* in an ontological sense. The *ξ* is a one-time event that has already happened under certain conditions and *ξ′* has to differ from *ξ* in some aspect. The best standard *ξ′* can purport to achieve is to capture relevant elements of *ξ* in a such way that performing inference about *R* while adhering to *A* and sampling the same population is possible within an acceptable margin of error. However, every experiment is embedded in its immediate social, historical, and scientific context, making it a non-trivial task for scientists to include all the relevant *K* when they report the experiment in an article and make explicit all the natural interpretations used to assign meaning to its results. As such, designing and conducting replication experiments cannot be reduced to a clerical implementation of reported experimental procedures. A comprehensive understanding of *K* is increasingly critical as *ξ′* diverges further away from *ξ* to be able to comprehend the nature and importance of the divergence for the interpretability of *ξ′* and for results reproducibility. For *ξ′* to serve their intended objective, information readily available from *ξ′* needs to be supplemented by a careful historical and contextual examination of the relevant literature and the broader scientific background. Otherwise, *ξ′* may differ from *ξ* in non-trivial ways impacting the meaning of the evidence obtained and changing the estimated reproducibility rate.

### 4.2 Model, *M*_*A*_

For *ξ′* to be able to reproduce *all possible R* of *ξ, M*_*A*_ must be specified up to the unknown quantities on which inference is desired. This specification must be transmitted to *ξ′*, such that *M*_*A*_ and 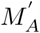 are identical for inferential purposes mapping to *R*. If an aspect of *M*_*A*_ that has an inferential value mapping to *R* is not transmitted to *ξ′* and this inferential value is lost, then *R* cannot be meaningfully reproduced by *R′*. On the other hand, given an inferential objective mapping to a specific *R*, the aspects of *M*_*A*_ that are irrelevant to that inferential objective need not be transmitted to *ξ′* to meaningfully reproduce *R* by *R′*. Counterintuitively, to meaningfully reproduce *R* by *R′, M*_*A*_ and 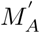 do not need to be identical, as given by result 3.

#### Result 3.

*M*_*A*_ and 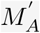 do not have to be identical in order to reproduce a result *R* by *R′*. Under mild assumptions, the requirement for *R* to be reproducible by *R′* is that there exists a one-to-one transformation between *M*_*A*_ and 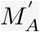 for inferential purposes mapping to *R* (proof and details in Appendix 4).

As an example of result 3, consider *ξ*_*bin*_ and *ξ*_*negbin*_ in figure 1. Conditional on the objective of estimating *p*, the population proportion of black ravens, any of (*ξ*_*bin*_, *ξ*_*bin*_), (*ξ*_*bin*_, *ξ*_*negbin*_), (*ξ*_*negbin*_, *ξ*_*bin*_), (*ξ*_*negbin*_, *ξ*_*negbin*_) can be effectively considered a pair (*ξ, ξ′*) of an idealized experiment and its (*exact*) replication. The reason is that the quantity of interest *p* is an identifiable parameter in both experiments although *M*_*A*_ and 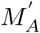 are not necessarily identical^4^.

In practice, when conducting a sequence of replication experiments, we would be interested in gauging the extent to which we can reproduce a specific result. Assuming that *S* are the same throughout all experiments, we expect the observed reproducibility rate of a sequence of experiments whose elements are chosen from *ξ*_*bin*_, *ξ*_*negbin*_ to converge on the same value, capturing the information on *p*, in the same way. However, result 3 does not imply that the (true) reproducibility rate of any two sequences of experiments involving any *M*_*A*_ and 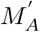 are equal to each other. In fact, the (true) reproducibility rates of two sequences are not equal, when non-exact replications are involved.

Openness of *M*_*A*_ to 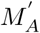 needs to be distinguished from the equivalence of *M*_*A*_ and 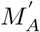. In *ξ*_*bin*_ and *ξ*_*negbin*_, 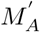 is not equivalent to *M*_*A*_. However, the binomial and the negative binomial models become equivalent with respect to a certain inferential objective that allows for reproducing a specific *R*, which is estimating *p*. To establish this compatibility, *M*_*A*_ should be open to *ξ′* but does not need to be assumed in *ξ′*. Specifically, to set 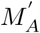 to be the negative binomial model in *ξ′* to reproduce the estimate of *p* in *ξ*, we need to know that *ξ* has used the binomial model. This ensures that *ξ′* can use a model that has the same parameter *p* with the exact same meaning as in *ξ* and same population assumptions *A* such that the inferential equivalence holds. A counterexample where *A* is different and this matters for reproducing a specific *R* is *ξ*_*hyper*_. *ξ*_*hyper*_ samples from an arbitrary finite subset of infinite population but still uses the same parameter *p* as *ξ*_*bin*_ and *ξ*_*negbin*_. The estimate of *p* in *ξ*_*hyper*_ will be biased due to differences in *A*. Without access to full specification of *M*_*A*_, this compatibility between *M*_*A*_ and 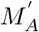 or lack thereof cannot be established.

This point is illustrated in many-analyst studies (Botvinik-Nezer et al., 2020; Silberzahn et al., 2018) in which a fixed *D* is independently analyzed by multiple research teams who are provided *D* and a research question that puts a restriction on which *R* would be relevant for the purposes of the project. The teams were not, however, provided a *M*_*A*_, *S*_*post*_, or full specification of *K*. Teams used a variety of models differing in their assumptions about the error variance and the number of covariates (*M*_*A*_) to analyze *D*. The results differed widely with regard to reported effect sizes and hypothesis tests. So even when *D* was open, the lack of specification with regard to *M*_*A*_ yielded largely inconsistent results. It is not because the same aspects of reality cannot be captured by different models but because researchers did not automatically agree on which aspects to capture in their models.

Taking stock, our ravens example is deliberately simple to help in our analysis. State of the art models are often large objects. If *M*_*A*_ is large, it might not always be clear which class of models 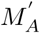 can be drawn from to be equivalent to *M*_*A*_, and finding this class might be unfeasible. Then *M*_*A*_ needs to be both open to and photocopied by *ξ′* to be able to reproduce the results of interest. This point is particularly important to communicate to scientists who primarily engage in routine null hypothesis significant testing procedures and may not be conventionally expected to transparently report their models^5^.

### 4.3 Method, *S*

#### 4.3.1 Pre-data methods, *S*_*pre*_

*S*_*pre*_ comprises a wide range of procedural components in *ξ* that feeds into collection of *D*_*v*_. Examples of *S*_*pre*_ are determining types of observables, unobservables, and constants, measurement and instrumentation choices, and sampling procedures such as random number generators used in computational methods.

Pertaining to mathematical features of the variables of interest, *S*_*pre*_ may capture their types or a particular scaling. For example, a variable can be assumed discrete, continuous, or both discrete and continuous for mathematical convenience. This choice determines whether we are bound by a counting measure or a Lebesgue measure. A variable can also be assumed categorical, ordinal, interval, or ratio. Some variables or parameters are scaled to the interval [0, 1] on the real line, to make their interpretation natural. All of these *S*_*pre*_ choices affect *M*_*A*_ and the consequent *S*_*post*_.

Pertaining to operational features of the variables of interest, Spre may capture the method of observation and measurement instruments. In our toy example, a raven can be observed for its color by naked eye (*S*_*pre*_), but another investigator may opt for a mechanical pigment test 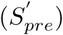. What considerations should be given when making substitutions for *S*_*pre*_? One issue due to choices in operationalization is measurement error. Measurement error in observables, when not accounted for, might be a factor unduly exacerbating irreproducibility or inflating reproducibility (Devezer et al., 2021; Loken and Gelman, 2017; Stanley and Spence, 2014). Another issue arises due to arbitrary choice of experimental manipulations or conditions which might not be mathematically equivalent. For example, manipulations that are not tested for specificity may end up manipulating non-focal constructs or only weakly manipulate the focal construct (i.e., leading to small effect sizes) (Gruijters, 2022).

Even though knowing all these features is useful in understanding *S*_*pre*_, there is a caveat. All aspects of *S*_*pre*_ must be fixed before realizing *D*_*v*_ and it is challenging to assess a priori whether *ξ* and *ξ′* using different *S*_*pre*_ and 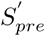 respectively can be equivalent to each other. Due to these complexities and ambiguities surrounding *S*_*pre*_, openness of *S*_*pre*_ seems to be the easiest way to obtain an equivalent 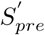 in designing and performing *ξ′*. However, there are well-known examples to show that *S*_*pre*_ and 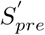 can be different and yet *ξ* and *ξ′* can be equivalent conditional on *R*, which leads us to result 4.

##### Result 4.

*S*_*pre*_ and 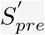 do not have to be identical in order to reproduce a result *R*.

As an example of result 4, consider models *ξ*_*poi*_ and *ξ*_*exp*_ in figure 1. *ξ*_*poi*_ has a good approximative model to the model in *ξ*_*bin*_ if we think of sampling ravens continuously from a population where black ravens are rare. We assume *np* → *λ*, where *λ* is rate of sampling the black ravens (parameter of the Poisson model) and under this assumption, we focus on inference on *λ*. Now, as a thought experiment, let us assume that we do not have a device to count the number of black ravens past 1. However, we have a chronometer. As a result of using the model in *ξ*_*poi*_, we are, as a mathematical fact, also using the model *ξ*_*exp*_, which measures the *time* between observing black ravens. Further, the two models have the same parameter, with the same interpretation. Therefore, if we were to measure the time between observing black ravens for a sample, then we can still perform inference on the rate of observing black ravens from the population. We note that *ξ*_*bin*_, *ξ*_*negin*_, *ξ*_*hyper*_, *ξ*_*poi*_, *ξ*_*nor*_ operate under different assumptions, but are still *counting* ravens and interested in the number of black ravens. In contrast, *ξ*_*exp*_ is considerably different from these experiments. It is *not* counting ravens, but *measuring time*, which we would reasonably define as a continuous variable. While *S*_*pre*_ in *ξ*_*exp*_ differs considerably from all other experiments in our toy example, the exponential experiment would serve as a meaningful *ξ′* to reproduce *R* in any of them, at least approximately.

#### 4.3.2 Statistical methods, *S*_*post*_

Statistical methods, *S*_*post*_, that are designed for a specific inferential goal, *R*, but do not return identical values when applied to a fixed *D* are common. Conversely, some statistical methods return identical values for a specific inferential goal, *R*, and they are mathematically equivalent conditional on *D*, even though they operate under distinct motivating principles. We have the following result.

##### Result 5.

*S*_*post*_ and 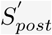 do not have to be identical in order to reproduce a result *R* by *R*^′^.

For the experiments *ξ*_*bin*_ and *ξ*_*negbin*_ in our toy example, the maximum likelihood estimator (MLE) and the method of moments estimator (MME) of *p* are numerically equivalent (see Appendix 5). This equivalence holds even when the interpretation of probability differs between methods. For example, MLE and the posterior mode in Bayesian inference under uniform prior distribution on parameters are equivalent regardless of all else.

At the minimum, for *ξ′* to be a meaningful replication of *ξ* conditional on *R*, the modes of inference should be equivalent. That is, the pair (*S*_*post*_, 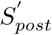) should belong to one of: point estimators, interval estimators, hypothesis tests, predictions, or model selection. Further, *S*_*post*_ should be open to *ξ′* but it does not need to be duplicated to establish equivalence. For example, to use MME to estimate *p* in *ξ′*, we need to know that *ξ* has used MLE or MME. This way, we can ensure that *ξ′* will at least use a numerically equivalent estimator as the one used in *ξ*, even if not equivalent in principle. On the other hand, it is well-known that a variety of *S*_*post*_ for the same mode of inference may yield different *R*. The many-analyst project by Silberzahn et al. (2018) provides clear examples of this. Teams which were given a fixed *D* to analyze for a pre-determined *R* (i.e., effect size as given by odds ratio), ended up implementing their choice of *S*_*post*_. Even when their modeling assumptions matched, the results they reported varied. For instance, out of the teams that assumed a logistic regression model with two covariates, one pursued a generalized linear mixed-effects model with a logit link for *S*_*post*_ (Silberzahn et al., 2018, *line 15 in Table 3) and another pursued a Bayesian logistic regression (Silberzahn et al*., *2018, line 16 in Table 3). The confidence intervals around the effect size estimates reported by these two teams do not even overlap despite using a fixed D*.

### 4.4 Data, *D*

#### 4.4.1 Data structure, *D*_*s*_

In statistics and philosophy of statistics, *D′* is often seen as the *new data* of the *old kind* in the sense that *D*_*v*_ and 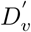 are independent of each other but *D*_*s*_ and 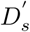 are identical. However, conditional on *R*, we have result 6.

##### Result 6.

*D*_*s*_ and 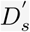 do not have to be identical in order to reproduce a result *R* by *R*^′^.

As an example of result 6, we consider the models in *ξ*_*poi*_ and *ξ*_*exp*_ in figure 1. Poisson model *counts* the black ravens as observable. It assumes that black ravens are observed with a constant rate. Exponential model measures the *time* between arrivals of black ravens. It also assumes that black ravens are observed with a constant rate. By referring to the unit of observations we see that the data structures in *ξ*_*poi*_ and *ξ*_*exp*_ are distinct. And yet, the unknown parameter about which inference is desired is the same, *λ*—the rate of black ravens appearing in continuous sampling (see Appendix 6).

As another example, note that the stopping rules of *ξ*_*bin*_ and *ξ*_*negbin*_ are different from each other. The stopping rule affects *D*_*s*_ because the maximum number of black ravens in *ξ*_*bin*_ is *n* but in *ξ*_*negbin*_ it is the maximum number of black ravens in the population. And yet, the estimate of *p* is the same in both experiments.

Data sharing is sometimes viewed as a prerequisite for a reproducible science (Hardwicke et al., 2018; Molloy, 2011; National Academies of Sciences, Engineering, and Medicine, 2017; Stodden, 2011). Our analysis suggests that this statement requires further qualification and calls for attention to *D*_*s*_. Result 6 notwithstanding, changes in *D*_*s*_ are not trivial and they impact the true reproducibility rate. For example, *ξ′* might be designed to have a larger sample size than that of *ξ*. In this case, the variance of the sampling distribution of the sample mean decreases linearly with the sample size and hence it would be different for *ξ* and *ξ′*. Typically, larger sample sizes are pursued to increase the statistical power of a hypothesis test in *ξ′*. While such *ξ′* will indeed increase the power of a test, it also impacts the reproducibility rate. Counterintuitively, under some scenarios this might play out as reproducing false results with increased frequency (see Devezer et al., 2021, for such counterintuitive results).

#### 4.4.2 Data values, *D*_*v*_

Having open access to *D*_*v*_ has no bearing on designing and performing a meaningful *ξ′*or on the reproducibility of *R*. Conditional on *R, ξ′* aims to reproduce *R*, not *D*_*v*_. Therefore, reporting *R* from *ξ* is sufficient for *ξ′* to assess whether *R* is reproduced by *R′*. However, information from *ξ* can be reported in a variety of ways and does not necessarily contain *R*. We show this with an example. We consider a model selection problem with three models *M*_1_, *M*_2_, *M*_3_, where *ξ* and *ξ′* use Some Information Criterion (SIC) as *S*_*post*_. Assume *ξ* reports selecting *M*_1_ as *R*. This is all *ξ′* needs to import to know whether *R* is reproduced in *R′*. If *R′* reports *M*_1_ as the selected model, then it is reproduced, else it is not. However, if which model is selected is not reported as *R, ξ′* needs values of SIC from *ξ* for all *M*_1_, *M*_2_, *M*_3_, so that *ξ′* can redo the analysis of *ξ* to find out what *R* was. In the unlikely event that not even SICs are reported, *ξ′* would need *D*_*v*_ to re-perform the whole analysis of *ξ* by applying *S*_*post*_ to *D* to calculate SICs and then get *R*.

##### Result 7.

*ξ* does not have to be *D*_*v*_-*Open* in order for *ξ′* to reproduce a result *R*.

That said, openness of *D*_*v*_ might facilitate auditing of *R* and vetting it for errors. There may be other benefits to open *D*_*v*_ such as enabling further research on *D*_*v*_ (e.g., meta-analyses). The distinction we draw matters particularly when there may be valid ethical concerns regarding data sharing (Borgman, 2012). Open *D*_*v*_ is best evaluated on its own merits as has been discussed extensively (Janssen et al., 2012) but cannot be meaningfully appraised as a facilitator of replication experiments or precursor of results reproducibility. While some level of open scientific practices is necessary to obtain reproducible results, open data are not a prerequisite.

## 5 Exact versus non-exact replications: A simulation study on reproducibility rate

So far we have established that to reproduce *R*, all elements of *ξ* do not need to be open, and not all elements that are required to be open need to be duplicated for a meaningful *ξ′*. On the flip side, we also established that relatively simple openness considerations such as experimental procedures, hypotheses, analyses, and data will not suffice to make *ξ′* meaningful. The challenge in making *π*-openness useful for replication experiments is to clearly identify and delineate the elements of the idealized experiment. For example, proper *K* is difficult to define and communicate with precision. Also, *M*_*A*_ is at times conflated with *S*_*post*_ and left opaque in reporting. As we discussed earlier, making *K* explicit and clearly specifying *M*_*A*_ up to its unknowns is critical when designing *ξ′*.

Hitherto, we focused on replication experiments and only alluded to results reproducibility when needed. In this tack, we have mathematically isolated *ξ* from *R*, and made some statements about *ξ* unconditional, and then conditional on *R* to emphasize their difference. Now that we turn our attention to explicitly drawing the link from replications to reproducibility, we condition *R* on *ξ*.

Given a sequence of *exact* replication experiments *ξ*^(1)^, *ξ*^(2)^, … and a result *R* from an original experiment *ξ*, do we expect to confirm *R* with high probability irrespective of the elements of *ξ*? The answer is “no” as shown elsewhere (Devezer et al., 2019, 2021). The true reproducibility rate of a result is a function of not only the true model generating the data but also the elements of the idealized experiment. *ξ* may be characterized by a misspecified *M*_*A*_ (e.g., omitted variables, incorrect formulation between variables and parameters), unreliable *S*_*pre*_ (e.g., measurement error, confounded designs, non-probability samples), unreliable *S*_*post*_ (e.g., inconsistent estimators, violated statistical assumptions), errors in *D* (e.g., recording errors), or large noise to signal ratio (e.g., large error variance and small expected value). All of these lead to the mathematical conclusion that the true reproducibility rate *ϕ* is specific to each configuration of *ξ* and thus can take any value on [0, 1]. Therefore, *ϕ* tells us more about the experiment itself than some unobserved reality that is presumed to exist beyond it. Since we are now conditioning on *ξ* and questioning the reproducibility rate of *R*, the conclusion is that while a degree of openness may be able to address a “replication” crisis by facilitating faithful replication experiments, it does not suffice to solve any alleged “reproducibility” crisis.

Openness of elements of *ξ* facilitates *ξ′*, thereby allowing us to estimate *ϕ* of *R* by *ϕ*_*N*_ conditional on *ξ*. However, *ϕ* cannot be reasonably used as a target of scientific practices where each *ξ* is designed to maximize it. It does not make sense to think that a *ξ* that returns the highest reproducibility rate for a given *R* is scientifically most relevant or most rigorous experiment. For example, choosing an *S*_*post*_ that always returns the same fixed value regardless of *D*_*v*_ would yield *ϕ* = 1. In fact, *ϕ* can be made independent of what it would be under sampling error^6^.

A reasonable expectation from *ξ′* is to deliver a scientifically relevant estimate of *ϕ*, given *R*. Openness plays an important role in this regard. In section 4 we established that any non-open elements of *ξ* would need to be substituted for in *ξ′*, leading to a non-exact replication. The following result states how a sequence of non-exact replications alter the reproducibility rate.

### Result 8.

Assume a sequence *ξ*^(1)^, *ξ*^(2)^, …, *ξ*^(*J*)^ of idealized experiments in which a result *R* is of interest. Then, the estimated reproducibility rate of *R* in this sequence converges to the mean reproducibility rate of *R* in *J* replication experiments. (See Appendix 7 for proof.)

Result 8 states that the true reproducibility rate to which the estimated reproducibility rate of a sequence of non-exact replication experiments converges is the mean reproducibility rate of results from all experiments in the non-exact sequence and not the true reproducibility rate of a fixed original result. Hence, the reproducibility rate is a function of all elements of the idealized experiment, for both a fixed original experiment and all its replications. Each replication that is non-exact in a different way from others introduces variability, decreasing the precision of estimates given a fixed number of replications.

We illustrate the link between replication experiments and reproducibility rate with a simulation study. We consider a series of exact and non-exact replication experiments to analyze the variation in the reproducibility rate of a result as a function of the elements of *ξ*. We use sequences of two idealized experiments *ξ*_*poi*_ and *ξ*_*nor*_, which are approximate models to binomial from our toy example. For all conditions, we fix the true proportion of black ravens and the number of trials in the exact binomial model at 0.01 and 1000, respectively. These arbitrary choices make the true reproducibility rate distinct under *ξ*_*poi*_ and *ξ*_*nor*_. As *R*, we choose a point estimate for the location parameter of the probability model. For convenience, we assume that the parameter estimates of the original experiments are equal to the true value. After each replication experiment we determine whether this result is reproduced by *R′* based on whether it falls within some suitably scaled population standard deviation units of the true parameter value.

In exact replications, we vary *M*_*A*_, *S*_*post*_, *D*_*s*_ of the idealized experiment, each element taking two values. This results in a 2 (*M*_*A*_) x 2 (*S*_*post*_) x 2 (*D*_*s*_) study design (8 conditions) for exact replications where: 1) Model assumed, *M*_*A*_ ∈ {*ξ*_*poi*_, *ξ*_*nor*_}, 2) Method as point estimate, *S*_*post*_ ∈ {MLE, posterior mode}, 3) Sample size, *D*_*s*_ ∈ {30, 200}. When *S*_*post*_ is the posterior mode, we use conjugate priors: Gamma distribution with rate and shape parameters 5 (arbitrarily chosen) for *ξ*_*poi*_, and Normal distribution with prior mean 10 and prior precision 1 for *ξ*_*nor*_. In figure 3, panels A and B show 100 independent runs of a sequence of 1000 exact replication experiments under these conditions, for *ξ*_*poi*_ and *ξ*_*nor*_, respectively.

**Figure 3.**
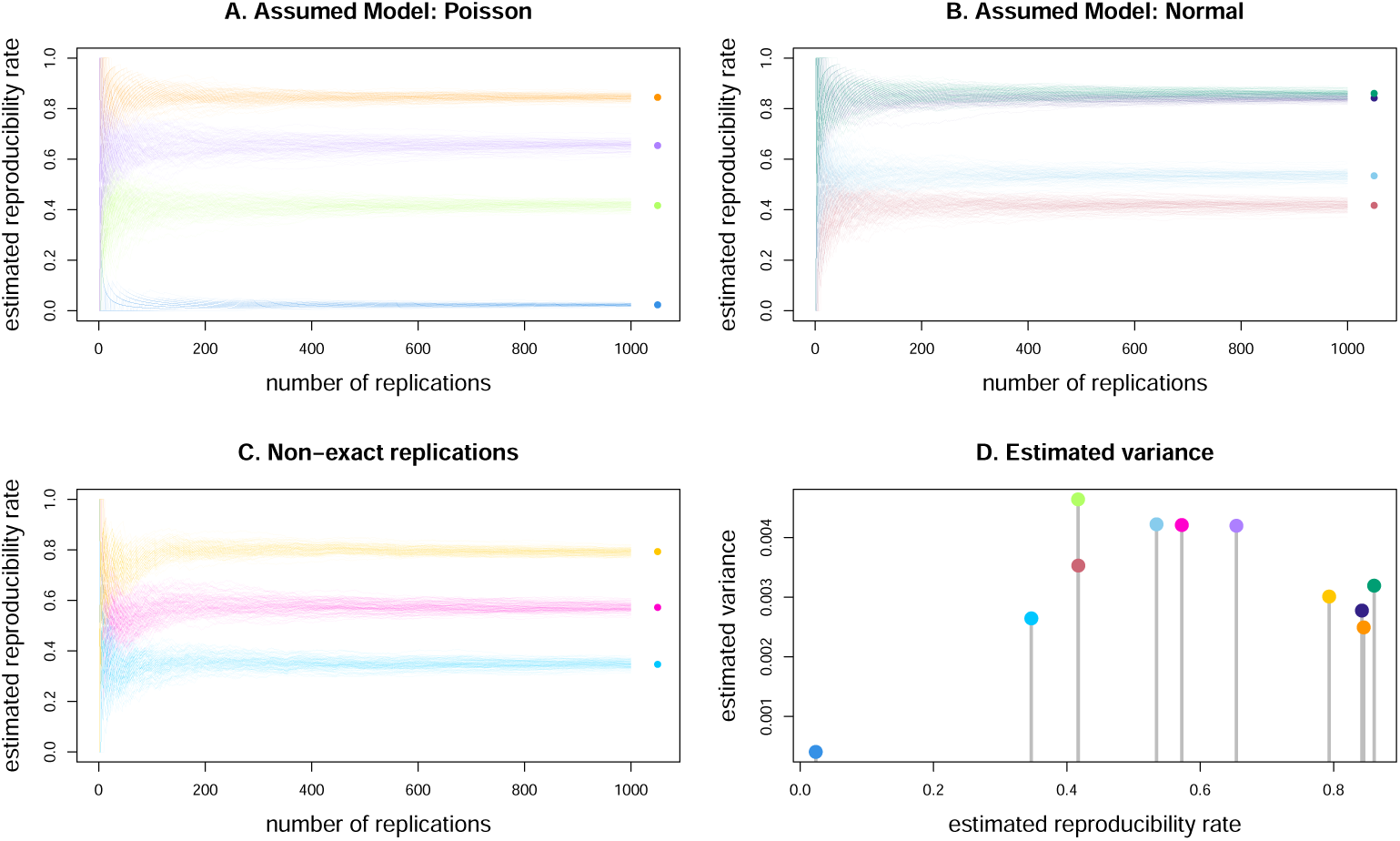
Reproducibility rates of a true result in sequences of 1000 exact (**A**. and **B**.) and non-exact (**C**.) replication experiments. *S*_*post*_ is varied as MLE and Posterior mode, and *D*_*s*_ is varied as *n* = 30 and *n* = 200. Each condition is color coded and consists of 100 independent runs. **A**. *M*_*A*_ : Poisson. 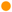 MLE, *n* = 200; 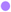 Posterior mode, *n* = 200; 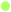 MLE, *n* = 30; 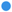 Posterior mode, *n* = 30. **B**. *M*_*A*_ : Normal. 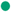 Posterior mode, *n* = 200; 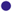 MLE, *n* = 200; 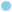 Posterior mode, *n* = 30; 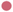 MLE, *n* = 30. **C**. Three cases of 1000 non-exact replication experiments where they are chosen uniformly randomly from the set of 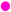 all eight idealized experiments, 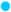 four idealized experiments with lowest reproducibility rates, 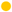 four idealized experiments with highest reproducibility rates. In **A**., **B**., **C**., * is the mean of the reproducibility rates of 100 runs at step 1000, an estimate of the true reproducibility rate for the sequence of idealized experiments. **D**. Variances of all 11 exact and non-exact sequences at step 50 of the simulation with respect to the estimated reproducibility rate (see text for interpretation).

In non-exact replications, we vary the set from which the replication experiment is uniformly randomly chosen from in each step. This results in additional 3 conditions: A set of all 8 idealized experiments, a set of 4 idealized experiments with lowest reproducibility rates, and a set of 4 idealized experiments with highest reproducibility rates. Panel C shows 100 independent runs of a sequence of 1000 non-exact replication experiments under these conditions.

We emphasize that all parameters of the simulation example in figure 3 are chosen so that the implications of differences between different models, methods, and data structures make the link between replications and reproducibility explicit. It is certainly possible to choose these parameters to obtain any true reproducibility rate defined by a specific *ξ* since *ϕ* ∈ [0, 1].

Conditional on *R*, some conclusions from figure 3 are as follows.

1. The true reproducibility rate depends on the true data generating mechanism and the elements of the original experiment. Specifically, the true reproducibility rate in our simulation is a function of the true model generating the data, *M*_*A*_, and also *D*_*s*_ such as the sample size, and *S*_*post*_ such as the method of point estimation. This can be seen from exact replication sequences of 8 idealized experiments in panels A and B, with the true reproducibility rate for each experiment indicated by stars.
2. By weak law of large numbers, even if the true reproducibility rate is high (e.g., orange in panel A and green in panel B), the estimated reproducibility rate from a short sequence of exact replications has higher variance than the variance of the estimated reproducibility rate in a longer sequence. However, the estimated reproducibility rate from exact replications ultimately converges to the true reproducibility rate of an original result from a fixed *ξ* illustrating result 1.
3. Estimated rate of reproducibility from a sequence of non-exact replications may be drastically different from the true reproducibility rate of an original result. The sequence of idealized experiments shown in pink in panel C of figure 3 is a sequence of non-exact replications for any of the 8 original idealized experiments in panels A and B. For example, assume the original experiment we aim to replicate is *ξ*_*poi*_ with *S*_*post*_ and *D*_*s*_ set to posterior mode and sample size *n* = 30, respectively. Blue sequences in panel A show that the true reproducibility rate of *R* (i.e., the estimate of location parameter) from these sequences of exact replication experiments is close to zero as shown by the convergence of 100 runs (i.e., blue star). If *S*_*post*_ and *D*_*s*_ were not open in this experiment then we would have had to substitute for them and the pink sequences in panel C would serve as plausible replication experiments. In this case, we would estimate the reproducibility rate of *R* as approximately 60% (i.e., pink star).
4. In a sequence of replication experiments, the set we choose the experiments from matters for true reproducibility rate. An original idealized experiment and its non-exact replications belonging to a set of idealized experiments that have true reproducibility rates close to each other for a given *R* yield an estimated reproducibility rate that is closer to the true value of the original experiment. For example, the yellow and blue sequences in panel C come from a set of 4 idealized experiments with the lowest and highest reproducibility rates among all 8 experiments, respectively. Compare the set of experiments sampled in the blue sequence to the set of experiments sampled in the yellow sequence. The latter serves as a more relevant set of idealized experiments for replications of the orange and purple experiments in panel A and the dark blue and green experiments in panel B, yielding a better reproducibility rate estimate for the original *R*. This pattern is an illustration of the broader theoretical result 8. In practice, however, we do not have access to the true reproducibility rate of any idealized experiment to help determine our replication sets. We have to make our decision based on the elements of the idealized experiment instead, and that requires a thorough understanding of how each element of the idealized experiment impacts the reproducibility rate in a given situation.
5. The variance of the estimated reproducibility rate of results in a sequence of non-exact replications can be higher or lower than the variance of the estimated reproducibility rate in a sequence of exact replications of the original experiment. The pattern of variances we observe in panel D is a direct consequence of *nϕ* following a binomial distribution and result 8. As a mathematical fact of the binomial distribution, its variance is maximum at *ϕ* = 0.5 and decreases as the probability of success, *ϕ*, gets closer to 0 or 1. Hence, we expect our estimates to vary greatly in a sequence of non-exact replication experiments with moderate true reproducibility rates. If a sequence of non-exact replications come from a homogeneous set of very high (or very low) true reproducibility rates, we expect our estimates to vary little. On the other hand, we expect highest variation in our estimates from exact replications if *ϕ* = 0.5 from the original experiment and from non-exact replications if they are highly heterogeneous in their true reproducibility rates.

In sum, the mere choice of the elements of *ξ* impacts both the level of the true reproducibility rate and the variance of the estimated reproducibility rate. Any divergence in *ξ′* may move the estimated reproducibility rate away from the true value for an original result and increase the variance of its estimates. In Appendix 8, we provide a broader example for result 8 in the context of linear regression models, under a model selection (rather than parameter estimation) scenario, where both true and false original results are considered. This simulation study demonstrates a similar pattern of results to those presented in figure 3. Combined, simulation results confirm that reproducibility rate can take any value on [0, 1] depending on the elements of *ξ* even when the original experiment indeed captures a true result, there is no scientific malpractice, and meaningful replication experiments can be performed to reproduce *R*.

## 6 Discussion

In this paper we focused on scientific experiment as the critical unit of analysis, formalizing the logical structure of experiments toward building a theory of reproducibility. We clarified what makes for a *meaningful* replication experiment even when an exact replication experiment is not possible and established how openness of different elements of the idealized experiment contribute to it. We distinguished between the ability of a replication experiment to reproduce a result and the true reproducibility rate for that result. We showed that theoretically it is not possible to justify a *desired level* of reproducibility rate in a given line of research and to reach a high level of reproducibility rate via eliminating malpractice, requiring open procedures or data, or performing replication experiments. Now we discuss key insights from our findings.

### 6.1 Reproducibility and the search for truth

A layperson understanding of reproducibility to the effect that “if we observe a natural phenomenon, we should be able to reproduce it and if we cannot reproduce it, our initial observation must have been a fluke” is exceedingly misleading. A statistical fact is that reproducibility is not simply a function of “truth”. This was illustrated in (Devezer et al., 2019) and proved in (Devezer et al., 2021): True results are not perfectly reproducible and perfectly reproducible results are not always true (see Appendix 9 for proof). *True reproducibility rate* of a result and the variability in its estimator are determined by many factors including but not limited to the true data generating mechanism: The degree of rigor of the original experiment as assessed by the extent to which its elements are individually reliable and internally compatible with each other, the degree to which replication experiments are faithful to the original and how any discrepancies impact the results, the degree of rigor of the replication experiment wherever it diverges from the original, and how we determine for a result to be reproduced. Factors such as effect size, sampling error, missing background knowledge, and model misspecification (Box, 1976; Dennis et al., 2019) could render true results difficult to reproduce.

As a useful reminder, sampling error might be masked by the choice of method and other elements of the idealized experiment. A false result could be 100% reproducible due to the choice of estimation method. Therefore, judgments of reproducibility cannot exclusively be used to make valid inference on the truth value of a given result (see also Bak-Coleman et al., 2022, for a computational model with a similar conclusion).

Even if some form of a perfect experiment that captures ground truth and its exact replications exist, it might take many epistemic iterations of theoretical, methodological, and empirical research to achieve them (see Chang, 2004, p. 45, for a detailed discussion on epistemic iteration). We cannot expect to skip the arduous iterative process of doing science and hope to arrive at a non-trivially reproducible science with procedural interventions. In most fields and stages of science, focusing on maximizing reproducibility seems like a fool’s errand. For meaningful scientific progress, at the minimum we should take care to properly analyze the elements of the original experiment to assess how they might impact the true reproducibility rate and analyze the discrepancies of replication experiment(s) from the original to gauge how our reproducibility estimates may vary from the true value of the original result’s reproducibility. In the course of “normal science” (borrowing terminology from Kuhn, 1970), reproducibility of a result is more likely to tell us something about the experiments that generated the result and its reproducibility rate estimates than the lawlikeness of some underlying phenomenon.

### 6.2 Defining reproducibility

One aspect of reproducibility that often gets overlooked: how we define and quantify a result and its reproducibility also determines the true reproducibility rate. For example, in a null-hypothesis significance test, we might call a “reject” decision in a replication experiment as a successfully reproduced result if the original experiment rejected the hypothesis. On the other hand, we might instead look at whether effect size estimate of the replication experiment falls within some fixed error around the point estimate from the original experiment. Everything else being equal, the true reproducibility rates are expected to be different between these two cases using different reproducibility criteria.

Our findings hold under mathematical definitions of a result (definition 3) and of reproducibility rate (definition 5). In the absence of such theoretical precision, we often resort to heuristic, common sense interpretations of terms. In Appendix 1 we present a detailed argument on why and how theoretical precision matters and provide an example of a plausible measure of reproducibility without desirable statistical properties. Such lax standards in definitions invite unwanted or strategic abuse of ambiguities when interpreting replication results when we have a limited understanding of what we should expect to observe. Our surprise at “failed” replication results or delight in “successful” ones may not be warranted and what we observe could simply be a theoretical limitation imposed by our definitions rather than a reflection of the true signal that presumably exists in nature. For an extreme example, consider the following: We might call a result as reproduced if the replication effect size estimate falls on the real line. That would trivially give us a 100% reproducibility rate.

Whenever we evaluate replications and estimate reproducibility, it is incumbent on us to understand how we define our results, how we determine reproducibility, and how our measures should be expected to behave under specific conditions.

### 6.3 Reproducibility and openness

Open practices in science have been intuitively proposed as a key to solving the issues surrounding reproducibility of scientific results. However, a formal framework to validate this intuition has been missing and is needed for a clear discussion of reproducibility. The notion of idealized experiment serves as a theoretical foundation for this purpose. Using this foundation, we have distinguished the concepts of replication and reproducibility, showing how openness is related to meaningful replications. We have also distinguished between two types of reproducibility (Appendix 3). Whether elements from one experiment carry over to a replication experiment is only relevant to epistemic–as opposed to in-principle–reproducibility. In practice, however, resource constraints determine the availability and transferability of information between experiments. A realistic framework needs to provide a refined sense of which elements of an experiment need to be open to reproduce a given result, as opposed to simply saying “all of it”.

We have identified different levels and layers of openness, and examined their implications. An experiment that is completely open in all elements does not necessarily lead to reproducible results and an experiment that does not open its data does not necessarily hinder replication experiments. Nevertheless, irreproducible results sometimes raise suspicion and discussions typically turn towards concerns regarding the transparency of research or validity of findings. These discussions are typically driven by heuristic thinking about replications. Such heuristics might not hold and can lead to erroneous inferences about research findings and researchers’ practices. To move the needle forward, we have provided a detailed evaluation of which elements of an experiment need to be made open relative to some objective, and which do not. For example, while necessary to audit the results of a given experiment, data sharing is not a prerequisite for performing replications or reproducing results (contrary to some suggestions, for example by National Academies of Sciences, Engineering, and Medicine, 2017), but other elements of an experiment are. On the other hand, reporting model details, such as modeling assumptions, model structure, and parameters, becomes critical for improving the accuracy of estimates of reproducibility. Notably, even in recent recommendations for improving transparency in reporting via practices such as preregistration, models are typically left out while transparency of hypotheses, methods, and study design are emphasized (Nosek et al., 2018; van’t Veer and Giner-Sorolla, 2016). Also noteworthy is that some degrees of openness is difficult to attain, such as fully open background knowledge, often causing practical constraints to limit our choices for replication experiments.

When critical elements of an original experiment are not open, replication researchers would be forced to introduce substitutions in their experimental designs. Such substitutions, as we have illustrated, characterize non-exact replications and will likely alter reproducibility rates in different directions, contributing to the challenge of interpreting replication results. Strong theoretical foundations and well-defined shared empirical paradigms in a given area of research could help generate meaningful substitutions whose downstream consequences on inference are well-understood.

### 6.4 Choosing non-exact replications

Assuming a sequence of perfectly repeatable experiments is a theoretical convenience—one that especially frequentist statistics enjoys greatly. In scientific practice we lack the luxury provided by this assumption. Exact replications are practically impossible. Understanding the implications of result 8 is crucial in this respect. It states that any sequence of non-exact replications converges to a true reproducibility rate. This rate may or may not be scientifically meaningful for a specific purpose. Especially for a sequence of non-exact replications, it is hard to find a scientifically meaningful interpretation of what the reproducibility rate shows, even when it is high.

A proper understanding of the elements of the original experiment needs to precede any replication design. And wherever divergences from the original experiment are inevitable, we should strive to theoretically match new design elements to the original ones if our objective is to reproduce an original result. When that is not possible, simulations varying the degree and nature of these divergences would inform us on their impact on the reproducibility rate and can provide guidance in designing non-exact replication experiments. A lack of theoretical understanding in this regard poses significant constraints on the interpretability of replication results.

In cases where the original experiment suffers from design issues that make results predictably less reproducible, it is advisable to iteratively work toward improving the configuration of the idealized experiment first before attempting any non-exact replications (Feest, 2019). If there is nothing there to revisit, we might be better off saving our scientific curiosity and resources for more fruitful avenues. In fact, there is room for major theoretical advancements on why and how to choose replications.

### 6.5 Reproducibility of a result versus accumulation of scientific evidence

We hope that advancing theoretical understanding of results reproducibility helps delineate how and why it is different from other quantities that aim to measure the accumulation of scientific evidence. The notion of reproducibility is unique in the sense that it is anchored on the results of an initial experiment. To the contrary, meta-analytic effect size estimates, for example, focus on an underlying true effect, after accounting for variation between studies being meta-analyzed while robustness tests aim to assess to what extent estimated quantities of interest are sensitive to changes in model specifications. It is a widespread interpretation that reproducibility also speaks to the reliability or validity of an underlying true effect and can reasonably be used as a measure of evidence accumulation. It should be clear by now that this is a misconception. Truth certainly plays a role in reproducibility of a given result but not (always) too loudly as reproducibility primarily captures patterns specific to the original experiment. A replication experiment in reference to an original result is a particular kind of an idealized experiment that has the capacity for achieving certain scientific objectives, such as confirming a theoretically precise prediction under well-specified conditions (that is, attempting to account for sampling error as a last source of uncertainty after everything else has already been accounted for) or estimating the reproducibility rate of a particular result of a given experiment. For other scientific objectives, such as to make an initial scientific discovery, to pinpoint the conditions under which a precise and reliable signal can be captured, to aggregate evidence for a theorized phenomenon, or to gauge the robustness or heterogeneity of an observed phenomenon across contexts, there are other idealized experiments better suited to the task than replications (Bak-Coleman et al., 2022; Feest, 2019) such as systematic exploratory experimentation (Steinle, 1997), metastudies (Baribault et al., 2018), multiverse analyses (Steegen et al., 2016), meta-analyses, and continuously cumulating meta-analyses (Fletcher, 2021). The fact that scientists still care to meticulously design their experiments to be informative and meaningful has more to do with other scientific values and objectives than reproducibility.

In a sense, accumulation of scientific evidence in support of a finding requires epistemic iterations and triangulation by independent approaches and methods to achieve specific scientific objectives (e.g., discovering a new phenomenon, explaining a mechanism, predicting a future observation). This process leads to gradually eliminating uncertainty and enhancing our confidence in our theories and observations. On the other hand, attempts at reproducing a given result in replications prioritize understanding and fine-tuning the logical structure of experiments, which we see as human data generation mechanisms. Proper appreciation of this aspect of reproducibility is capable of guiding us in the right direction in our struggle to design more rigorous and informative experiments under uncertainty.

### 6.6 Concluding remarks

The discourse on scientific reform and metascience has so far pursued a “crisis” framing, focusing on behavioral, social, institutional, and ethical failings of the scientific endeavor and calling for immediate institutional and collective action. Our analysis shows that neither elimination of scientific malpractice nor actively encouraging replication experiments would necessarily improve the reproducibility of results. Because irreproducibility, when formally defined, appears to be an inherent property of the scientific process rather than a meaningful scientific objective to pursue. While reproducibility rate is a parameter of the system and thereby a function of truth, that view of the concept misses the big picture—that reproducibility reflects the properties of experiments. We perceive two issues with advancing a replication/reproducibility crisis narrative:

1. Conflating replication and reproducibility creates an inaccurate impression that these two alleged issues of not being able to conduct informative replication experiments and not being able to reproduce results are indistinguishable issues that can be addressed via similar solutions.
2. Framing irreproducibility as a crisis implies that there is an ideal rate of reproducibility we should expect or strive to achieve in a given field at a given time and we are falling short of this ideal standard.

Our mathematical results firmly argue against both of these misconceptions.

Shifting the discourse on scientific reform and metascience toward greater theoretical may help change the course of science. Instead of prioritizing crisis management measures, progress can be made by falling back on fundamental issues and working our way from the bottom up. That may require individual scientists to take a step back and reassess the way they have been practicing science. Circling back to our original premise, we emphasize that the problem is conceptual: The logical structure of experiments is not well understood and how experiments relate to reality gets misconstrued. Experiments are data generating machines and each element outlined in this work determines what kind of data they will generate. Gaining clarity with regard to how experiments impact the observed reality and properly assessing the empirical value of a given experiment for a given objective should precede concerns regarding possible replications. Theory of reproducibility is a step in this direction.

## Appendix 1

### Proof of result 1. And an example pertaining remark 2 that meaningful continuous measure of reproducibility which is nonetheless pathological

Result 1 is a consequence of Strong Law of Large Numbers. An easy proof relies on Kolmogorov’s almost everywhere convergence which states that a sequence of independently and identically distributed random variables with finite mean converges almost surely to a constant if and only if that constant is the expected value of random variables. The sequence *ϕ*^(1)^, *ϕ*^(2)^, …, *ϕ*^(*N*)^ obtained from *ξ*^(1)^, *ξ*^(2)^, …, *ξ*^(*N*)^ (respectively) satisfies Kolmogorov’s. By definition 5 *ϕ*_*N*_ ∈ [0, 1] and *ϕ*_*i*_ are independent of each other and identically distributed and the expected value is *E*(*ϕ*_*N*_) = *ϕ <* ∞, proving result 1.

Importantly, remark 2 cautions us that result 1 does not hold for all measures of reproducibility. A well defined *ξ* and *ϕ* are prerequisities for result 1 to hold. We use a counterexample with a continuous measure of reproducibility to clarify this point. As opposed to a 0-1 measure such as *ϕ*_*N*_, we consider a (maybe) more desirable measure of reproducibility rate, perhaps a degree of agreement between the results of *ξ* and *ξ′* to assess whether *r*_*o*_ from *ξ* is reproduced in *ξ′*. One way to represent this degree of agreement is to replace the indicator function in definition 5 with a function of a continuous random variable. For example, for a sequence of idealized experiments *ξ*^(1)^, *ξ*^(2)^, … we might define *Y* ^(*i*+1)^*/Y* ^(*i*)^, where *Y* ^(*i*)^ ∼ Nor(0, *σ*) is a centralized statistic from *ξ*^(*i*)^, as score on how extreme is a specific result with respect to an original result *Y* ^(*o*)^. Here *Y* ^(*i*)^ are independent and identically distributed random variables conditional on *ξ*^(*i*)^. The setup is such that if *Y* ^(*i*+1)^*/Y* ^(*i*)^ = 1, then the results in *ξ*^(*i*+1)^ and *ξ*^(*i*)^ have exactly the same degree of agreement. Thus, one can define the reproducibility rate as

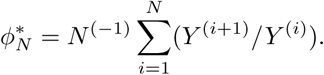

This measure of reproducibility rate might seem reasonable, but it is statistically unacceptable. To see this, we substitute *ϕ*_*N*_ with 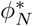 and we see that equation 1 is not true and we do not have desirable statistical properties for our estimator of reproducibility (Serfling, 1980, p.12). Consequently, the statistical justification for the concept of result reproducibility falls apart. This example shows that one has to define the parameter and its estimator of the reproducibility rate by obeying the constraints of statistically desired properties for reproducibility rate to be a useful concept. It is wise to check that a new concept defined in a developing field is statistically well-behaved. Statistical nuances might get lost in applications with important consequences for results reproducibility.

Some additional statistical properties of *ϕ*_*N*_ given in definition 5 are as follows. The sampling distribution of *ϕ*_*N*_ is asymptotically normal with *E*(*ϕ*_*N*_) = *Nϕ* and *V ar*(*ϕ*_*N*_) = *Nϕ*(1 − *ϕ*) by the Central Limit Theorem. All else being equal, the results for which the true reproducibility rate is high or low have low variance for the estimator and for the results for which the true reproducibility rate is around 0.5 the variance of the point estimator is large (largest when *p* = 0.5). Approximately 100% Confidence Intervals (and tests of approximately power 1) can arbitrarily be built, with the property that only finitely many of the confidence intervals do not contain the true reproducibility rate *ϕ*. This result, which fundamentally relies on the law of the iterated logarithm, constitute a strong basis for statistical methods about *ϕ*.

## Appendix 2

### Proof of result 2 (constructive): The sequence of idealized experiments *ξ*^(1)^, *ξ*^(2)^, … given by definition 5 is a proper stochastic process, seen as a joint function of random sample *D* and of each value in the support of data generating mechanism, *x* ∈ ℝ

*K, S, M*_*A*_ are not stochastic, so we condition on them. *ξ*^(*i*)^ draws a simple random sample *D*^(*i*)^ = **X**_**n**_^(*i*)^ independent of all else. We note two facts for the proof:

i. For fixed **X**_**n**_, the sample estimate of *M*_*A*_, is a well-defined probability model for all *x*. This setup induces the set of proper probability distribution functions: right continuous cumulative distribution functions with left limits on [0, 1]. Three of these cumulative distribution functions are exemplified in figure 4, left and middle panels.
ii. For any fixed *x* in the support of the cumulative distribution function of *M*_*A*_, the sample estimate of *M*_*A*_ as a function of the random data **X**_**n**_ is a random variable, which makes *ξ* a random variable. This is exemplified in figure 4, right panel, conditional on red line.

Together, (i.) and (ii.) imply that as a joint function of random *D* and *x, ξ* is a proper stochastic process (Serfling, 1980, Chapter 1-3) on the space of right continuous functions with left limits on [0, 1]. Examples are all gray cumulative distribution functions depicted in figure 4, right panel.

**Figure 4.**
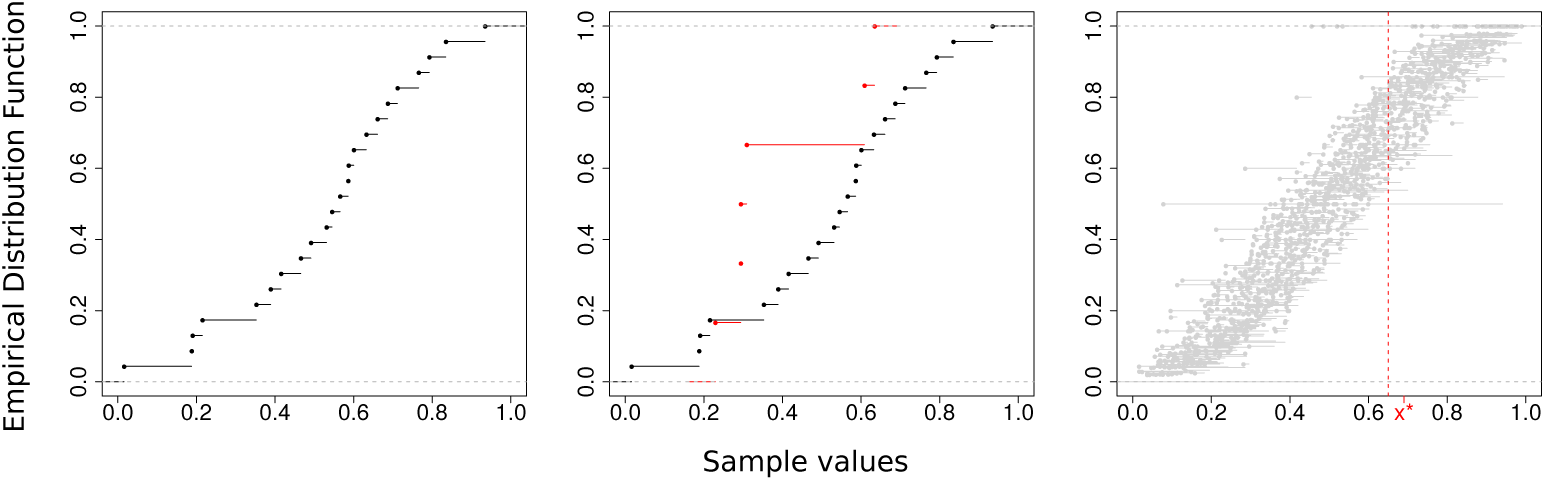
Left: Empirical CDF (ECDF) of a sample of size 30, emphasizing that the ECDF is a right continuous function. Middle: ECDF of the sample in the left panel (black) and that of an independent sample of size 10 (red) emphasizing that the ECDF is a random variable whose probability distribution is determined by the sample values (and hence data generating mechanism). Right: 100 independent samples of varying sample size (gray) emphasizing that ECDF is a stochastic process. Red vertical line shows the distribution of ECDF conditional on value *x*^*^.

Result 2 is a convenient way to study replications and reproducibility. It has a number of mathematical implications. First, it established that *ξ* is a well-behaved stochastic process with a limiting distribution. It is of interest to know the limit of this process. It tells us to which point the sample reproducibility rate from replication experiments converge.

Technically, the sequence of probability measures defined for the stochastic process associated with *ξ*^(1)^, *ξ*^(2)^, … on Borel sets with respect to the metric that we describe below has a limiting process that convergences in distribution. Establishing this convergence helps us to understand the limiting behavior of *ξ*^(1)^, *ξ*^(2)^, …, and characterizing this limiting behavior. Donsker’s Theorem characterizes the limiting process and states that *ξ* must convergence to the Wiener measure. Thus, the probability distribution of the reproducibility rate converges to the normal distribution. Readers interested in the theory of convergence in stochastic processes may refer to (Serfling, 1980, Chapter 1-3) for details. We give a brief description of necessary background here. There are three essential elements to study the convergence of a proper stochastic process: 1) A proper field on which the process takes values (the class of sets of interest) and a metric associated with it to assess the convergence of the process, 2) The probability measure that determines the behavior of the process, 3) Using (1) and (2), a complete mathematical formulation of the stochastic process which can be used to show convergence to some well-defined distribution.

We now consider a stochastic process as a function of *t* ∈ [0, 1], a random point in the space of right continuous functions on [0, 1] with left hand limits. We let the supremum of the L1 norm between any two points in the space and the metric to assess the convergence to be the classical Kolmogorov-Smirnov distance. By ⌊*nt*⌋ we denote the floor function, the integer part of *nt*. Given {**X**_**n**_ = (*X*_1_, *X*_2_, …, *X*_*n*_); *n* ∈ ℤ^+^}, where *X*_*i*_ are independent of each other and identically distributed, we define the stochastic process defined on partial sums:

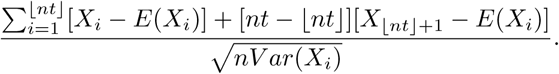

For elements of this process, if we denote the probability distribution for a sample size *n* by *P*_*n*_, then the limiting distribution is the well-known Wiener measure, 𝒲. Some results follow from this.

*ξ* is most generic when *M*_*A*_ is *any* probability model. This induces *S*_*post*_ having the sampling distribution function of *any* statistic. In this most generic case, the distribution of the sample reproducibility rate *ϕ*_*N*_ for the sequence *ξ*^(1)^, *ξ*^(2)^, … is asymptotically normal. To see this, we first let 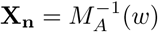, where *w* ∈ [0, 1] so that we have the image of the statistical model and assume that *ϕ*_*N*_ evaluated at 0 and 1 is 0. The stochastic process

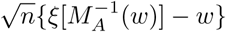

converges to a specific Wiener process, with bound end points, which is a Brownian Bridge: The process is Gaussian with zero expectation and for two points *w*_1_, *w*_2_ the covariance function *Cov*(𝒲 (*w*_1_), 𝒲 (*w*_2_)) = *w*_1_(1 − *w*_2_), with the ordering *w*_1_ ≤ *w*_2_, and *w*_*i*_ ∈ [0, 1].

By definition of this stochastic process and its convergence to a Brownian Bridge, we see that for each fixed value of *x, ξ* is asymptotically normally distributed with mean *M*_*A*_ and variance *M*_*A*_(1 − *M*_*A*_)*/n*.

The result can also be studied fixing one dimension at a time, giving two corollaries. For random data **X**_**n**_ the elements of the sequence of replication experiments *ξ*^(1)^, *ξ*^(2)^, … are random variables and conditionally independent of each other. For fixed data, the elements of the sequence of replication experiments *ξ*^(1)^, *ξ*^(2)^, … are probability models.

## Appendix 3

### Details on remark 2: Let *ξ* be an idealized experiment and *ξ′* be its exact replication. Conditional on *R* from *ξ, K′* is necessarily distinct from *K* for epistemic reproducibility of *R* by *R′*, but not necessarily distinct for in-principle reproducibility of *R* by *R′*

We define and distinguish *in-principle* reproducibility and *epistemic* reproducibility conditional on a result *R*. It is clear that *π*-openness where *π* is a non-empty set is necessary to make the elements of *ξ* available for replication *ξ′*. Further, *R* also needs to be open for *ξ′* to be able to determine whether *R′* has epistemically reproduced *R*. So, information on *R* across the sequence of replication experiments is a logical necessity for *epistemic* reproducibility. As an example, consider two scenarios 1 and 2. In each scenario, there are two experiments, the originals (*ξ*_1_ and *ξ*_2_, respectively) and their replications (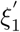 and 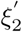, respectively). Each experiment assumes an infinite population of black and white ravens (*A*). *ξ*_1_ and *ξ*_2_ have identical *M*_*A*_, *S, D*_*s*_. *R* is the estimate 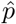 of the population proportion of black ravens *p*, obtained using an independent *D*_*v*_. We assume that the number of black ravens *b* observed in *ξ*_1_ and 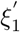, and *ξ*_2_ and 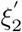 are the same.

#### Closed scenario

The experiments are isolated from each other and there is no information flow from *ξ*_1_ to 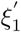. Thus, 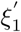 can only match all the elements of *ξ*_1_ that are relevant to 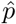 either by *chance* or by an extreme precision of prior theoretical formulation. By our example, *ξ*_1_ and 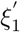 have identical *M*_*A*_, *D*_*s*_, *S* and have the same observed value *b* in the sample, thus they return the same estimate 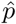. However, 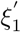 does not have any information pertaining *R* from *ξ*_1_, and thus 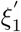 is in a position neither to learn from *R* of *ξ*_1_, nor to claim that it reproduced the result of *ξ*_1_ by *R′*. If an external observer were to observe the experiments *ξ*_1_ and 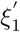, they could learn from the results of both experiments simultaneously. Starting with a prior view of equal proportion of black and white ravens, they could use the number of ravens observed in *ξ*_1_ and 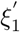, to conclude that *R* of *ξ*_1_ is indeed reproduced by *R′* of 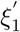 and arrive at an updated view. When there is no information exchange with regard to *R* between the *ξ*_1_ and 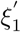, however, there is no meaningful or immediate *epistemic* interaction between *ξ*_1_ and 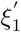, and there is no knowledge of reproducibility unless an all-knowing third party is involved.

This closed scenario shows that if there is no openness in the sense of information flow from one experiment to the next, it is improbable (but still possible) for an experiment to reproduce the result of another experiment. In order to acknowledge this point, we say that a result can only be *in-principle reproducible* if there is no epistemic exchange between *ξ*_1_ and 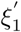 which could speak to the reproducibility of *R*, with the exception of via some omniscient external observer. At times historians of science illustrate such examples of scientific discoveries independently arrived at by different scientists unaware of each other’s work.

#### Open scenario

There is information flow from *ξ*_2_ to 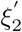, with respect to *R* and other information relevant to obtain 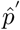 in 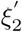. If 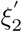 incorporates this information, it is a replication. Here, 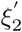 matches the elements of *ξ*_2_ by *social learning*. The information necessary for learning is transmitted in *K* and *R*. Starting with 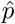 as 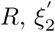 could conclude that they have indeed reproduced it. Thus, in the *open scenario* there is an *epistemic* interaction between *ξ*_2_ and 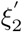 which contributes to the progress of science through deliberate transfer of knowledge via social learning, which gives us the notion of *epistemic reproducibility*.

As an example, we show the difference between *epistemic reproducibility* and *in-principle reproducibility* in figure 5 with an infinite population of black and white ravens and Bayesian inference. The panel on the left illustrates the closed scenario: Researchers of *ξ*_1_ assume a prior view of 1*/*2 on 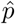. After observing *n* = 2 black ravens, they update their view to 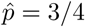 by Bayesian inference. Researchers of 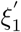 assume a prior view of 1*/*2 on 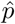 and observe identical *D*_*v*_, *n* = 2 black ravens as in *ξ*_1_, and they update their view with same *S*_*post*_, to reach 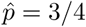. However, in the absence of an external observer these two results cannot be epistemically connected, thus reproducibility is only *in principle* in the absence of an external observer privy to both experiments. The panel on the right illustrates the open scenario: Researchers of *ξ*_2_ assume a prior view of 1*/*2 on 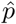. After observing *n* = 2 black ravens, they update their view to 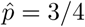 by Bayesian inference. 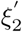 is a proper replication experiment. It is informed by the result of *ξ*_2_ as well as *K, M*_*A*_, *S, D*_*s*_ and observes identical *D*_*v*_ as 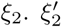, starting with a view of 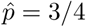 from *ξ*_2_, they update their view to 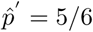. Thus, 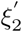 learns from *ξ*_2_, here in a Bayesian manner. The two results can be connected and thus reproducibility is *epistemic*.

**Figure 5.**
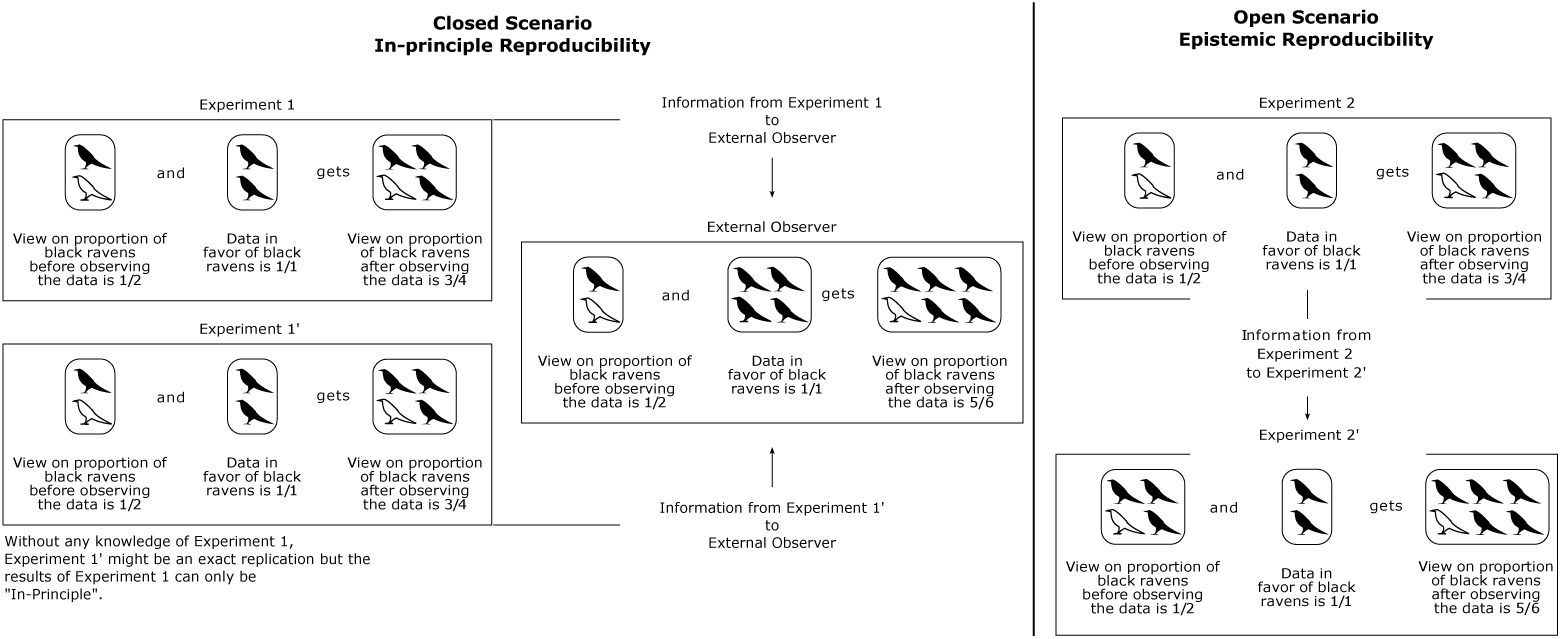
Epistemic versus in-principle reproducibility with an example of Bayesian information flow and learning (details within appendix text).

## Appendix 4

### Proof of result 3: *M*_*A*_ and 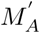 do not have to be identical in order to reproduce a result *R* by *R′*. Under mild assumptions, the requirement for *R* to be reproducible by *R′* is that there exists a one-to-one transformation between *M*_*A*_ and 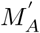 for inferential purposes mapping to *R*

We first give a proof for the statement and then follow with a specific example. We let *F*_*X*_(*x*) and *F*_*X*_*′* (*x*) be distribution functions with inverses 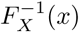 and 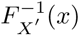, under *ξ* and *ξ*′, respectively. By assumption, a one-to-one function, *g*, from *F*_*X*_*′* (*x*) to *F*_*X*_′ (*x*) exists. For two distribution functions whose inverses exist, the mapping of population quantiles from one to the other also exists if there is a one-to-one function between these distribution functions. All well-behaved (non-order) statistics can be represented as quantiles, so we prove result 3 without loss of generality by setting the quantity of inferential interest as *x*_*q*_ where *F*_*X*_ (*x*_*q*_) = *P* (*X* ≤ *x*_*q*_) = *q* ∈ [0, 1]. If using equivalent estimators of quantiles with samples from *M*_*A*_ and 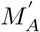 respectively, then the mapping carries over to *R* to *R′*. We have

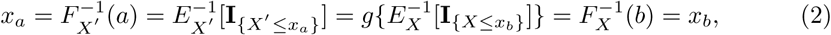

where **I**_{*A*}_ = 1 if *A* and 0 otherwise. Equations 2 hold for estimators of population quantities and an estimator of *x*_*a*_ can be equated to an estimator of *x*_*b*_ via a one-to-one transformation *g*, by replacing the population quantities with their estimators 2. This result applies to non-parametric and parametric models alike, in fact to all distributions with well-defined inverses.

As an example with two parametric models from figure 1 we consider the problem of estimating the proportion of black ravens, *p* using *ξ*_*bin*_ as the original experiment and *ξ*_*negbin*_ as its replication. The characteristic function of *ξ*_*bin*_ and *ξ*_*negbin*_ are (1 − *p* + *peit*)^*n*^ and *p*^*w*^(1 − *e*^*it*^ + *peit*)^−*w*^, respectively. The characteristic function for a random variable fully defines its probability model and thus, *ξ*_*bin*_ and *ξ*_*negbin*_ have distinct models. Yet *p* is an identifiable and estimable parameter of both experiments. The maximum likelihood estimator of *p* is 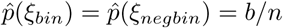 because *ξ*_*bin*_ and *ξ*_*negbin*_ are in a *likelihood equivalence class* with respect to parameter *p*. To see this, we note that the maximum likelihood estimator is obtained by setting the expression resulting from taking the derivative of the logarithm of the likelihood function (i.e, score function) with respect to *p* and solving for *p*. Under *ξ*_*bin*_ the score function is

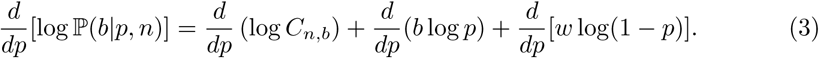

Under *ξ*_*negbin*_ the score function is

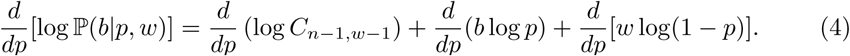

Equations 3 and 4 differ only in their first terms which is irrelevant to estimate *p* and thus 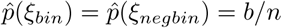 is the unique solution. The first terms on the right hand side of these two equations determine the stopping rule of the experiments. In *ξ*_*bin*_ we stop the experiment when *n* ravens are observed and the last raven can be black or white. In *ξ*_*negbin*_ we stop the experiment when *w* white ravens are observed and the last observation must be a white raven. This difference between stopping rules means that: 1) *S*_*pre*_ is different from 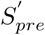. 2) Under our choice of *S*_*post*_ and 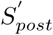 as the maximum likelihood estimator, the stopping rules in two models are irrelevant for estimating the proportion of black ravens in the population.

## Appendix 5

### Proof of result 5: *S*_*post*_ and 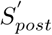 do not have to be identical in order to reproduce a result *R* by *R*^′^

There are a few heuristic ways to derive well-behaved statistical estimators of parameters. Examples include: method of moments, maximum likelihood, posterior mode (Bayesian). Well-known estimators may be equal to each other in value but motivated by distinct principles. For example, for some distinct probability models in the exponential family, the method of moments and the maximum likelihood estimator return the same value. Or, using uniform prior in Bayesian inference, the posterior mode always returns the same value as the maximum likelihood estimator. This motivates result 5 in the sense that *S*_*post*_ and 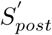 do not have to be identical to reproduce *R* by *R*^′^.

As an example based on *ξ*_*bin*_ from figure 1 we consider the following three estimators:

- If *S*_*post*_ is the maximum likelihood estimator motivated by the likelihood principle, then we have (see Appendix 4)

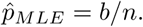
- If *S*_*post*_ is the method of moments estimator, the motivation is to set the population mean equal to the sample mean and solve for *p* and we have

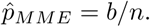
- If *S*_*post*_ is the posterior mode under the uniform prior (a special case of conjugate prior for *ξ*_*bin*_) we have

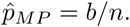

Therefore, *ξ* can employ any one of these three estimators as *S*_*post*_ and *ξ*^′^ can employ another as 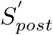 and still reproduce *R* by *R*^′^, as if they have used the same statistical method. For other modes of statistical inference such as hypothesis tests and prediction, we can find examples of numerically equivalent methods that are not identical in motivation (e.g., Shively and Walker, 2013).

## Appendix 6

### Proof of result 6: *D*_*s*_ and 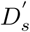 do not have to be identical in order to reproduce a result *R* by *R′*

The data structures of probability models that correspond to *ξ*_*bin*_, *ξ*_*negbin*_, *ξ*_*hyper*_, *ξ*_*poi*_, *ξ*_*exp*_, *ξ*_*nor*_ are all distinct. In *ξ*_*bin*_ and *ξ*_*negbin*_ the data structures are a sample of size *n* ravens and a sample of size *w* white ravens, respectively, from an infinite population in which *p* is constant. Stopping rules of the sampling in these experiments are different from each other: The last raven must be white in *ξ*_*negbin*_ but not in *ξ*_*bin*_. In *ξ*_*hyper*_, the stopping rule is the same as *ξ*_*bin*_, but the parameter *p* changes with each sample obtained due to finite population assumption in *ξ*_*hyper*_.

In *ξ*_*poi*_ and *ξ*_*exp*_, *np* → *λ* is the rate of black ravens appearing in the process. The observable in *ξ*_*poi*_ is the random variable *b*_*t*_, the count of black ravens at time *t* and we denote the count of black ravens at time *t* + *δ* by *b*_*t*+*δ*_. The observable in *ξ*_*exp*_ is the random waiting time *δ* to observe another black raven assuming a black raven is observed at time *t*. The equivalence between the parameters of *ξ*_*poi*_ and *ξ*_*exp*_ is given by

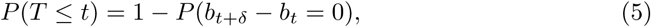

where the cumulative distribution function of the time variable in *M*_*A*_ in *ξ*_*exp*_ is related to the counts in *M*_*A*_ in *ξ*_*poi*_ by probability of no event in time period *δ*. Equation 5 implies that no black raven is observed in *δ*. By Poisson probability mass function we have this probability as *P* (*b*_*t*+*δ*_ − *b*_*t*_ = 0) = *e*^−*δ*^ and we have *P* (*T* ≤ *t*) = 1 − *e*^−*δ*^. This identifies *T* as an exponential random variable in *ξ*_*exp*_ implying that the data structures in *ξ*_*poi*_ and *ξ*_*exp*_ are distinct. Yet, irrespective of all these differences in data structures, *ξ*_*bin*_ and *ξ*_*negbin*_ estimate the same parameter, *p*. Further, *ξ*_*poi*_ and *ξ*_*exp*_ also estimate the same parameter, *λ*. Hence, *R* can be reproduced by *R*^′^ without the data structures being identical in *ξ* and *ξ′*.

## Appendix 7

### Proof of result 8: Assume a sequence *ξ*^(1)^, *ξ*^(2)^, …, *ξ*^(*J*)^ of idealized experiments in which a result *R* is of interest. Then, the estimated reproducibility rate of *R* in this sequence converges to the mean reproducibility rate of *R* in *J* replication experiments

Conditional on all other elements of an idealized experiment, definition 5 and consequently equation (1) assume that data are generated independently in each replication which implies that *R*^(*i*)^ are independent and identically distributed random variables. Result 8 is straightforward for independent and identically distributed random variables. Unconditionally on the elements, however, *R*^(1)^, *R*^(2)^, … in the sequence *ξ*^(1)^, *ξ*^(2)^, … are *not* independent and identically distributed, implying that *R*^(*i*)^ are not drawn from the same sampling distribution of results. An easy way to see this is to pick *ξ*^(*i*)^ and *ξ*^(*j*)^ distinct at least with respect to one element. In exact replications, *R*^(*i*)^ and *R*^(*j*)^ will converge to their unique true reproducibility rate *ϕ*^(*i*)^ and *ϕ*^(*j*)^ by equation (1). However, equation 1 can be generalized to obtain result 8 as follows using a theorem due to Kolmogorov (See Rao, 1973).

We let 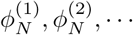 be estimates of reproducibility rates, with means *ϕ*^(1)^, *ϕ*^(2)^, … and variances *N* ^−1^*ϕ*^(1)^(1 − *ϕ*^(1)^), *N* ^−1^*ϕ*^(2)^(1 − *ϕ*^(2)^), …, respectively. We assume that the series 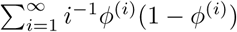 converges. Then,

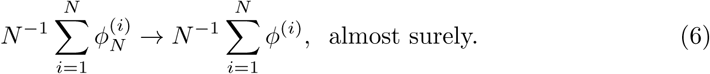

Expression (6) states that the estimated reproducibility rate of results from non-exact replication experiments meaningfully converges to the mean true reproducibility rate of the idealized experiments performed. The case of exact replications given by equation (1) is a special case of the equation (6), where all non-exact replications are identical to each other (and thus exact) with respect to the result obtained in an original idealized experiment. That is, if equation (6) is applied to *ξ* ≡ *ξ*^1)^ ≡ *ξ*^(2)^ ≡ … ≡ *ξ*^(*N*)^, where the true reproducibility rate for *R*_*o*_ obtained from *ξ* is *ϕ*, and we get

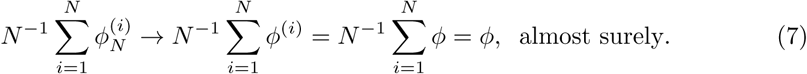

## Appendix 8

### Reproducibility rate of *R* as a model selection problem, in the context of linear regression models

In addition to the simulation example given in 3, here we present a second simulation example to illustrate the convergence of reproducibility rates from exact and non-exact replication experiments to their true value. Our example involves model selection problem in the context of linear regression models. Briefly, we assume the linear regression model

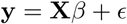

where **y** is *n* × 1 vector of responses, **X** is *n* × *k* matrix of fixed observables with first column entries equal to 1, *β* is *k* × 1 vector of parameters, and *ϵ* is *n* × 1 vector of independent and identically distributed normal errors with mean 0 and unknown variance. The statistical problem is as follows: Given *D* with *D*_*v*_ independent and identically distributed and *D*_*s*_ constituting *n* × 1 responses and *n* × *k* observables, select the best linear regression model among three models with respect to a model selection criterion (*S*_*post*_). The saturated model is given by

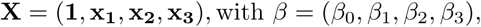

where **x**_**1**_, **x**_**2**_, **x**_**3**_ are *n* × 1 vectors of first, second, and third predictors,and *β*_1_, *β*_2_, *β*_3_ their respective regression coefficients. The set of three models considered in the model selection problem are:

1. **X** = (**1, x**_**1**_, **x**_**2**_, **x**_**3**_), with *β* = (*β*_0_, *β*_1_, *β*_2_, *β*_3_),
2. **X** = (**1, x**_**1**_, **x**_**2**_), with *β* = (*β*_0_, *β*_1_, *β*_2_),
3. **X** = (**1, x**_**1**_, **x**_**3**_), with *β* = (*β*_0_, *β*_1_, *β*_3_).

In all cases the true model generating the data is model 3. For each *ξ* (and their exact replications), we vary four elements *M*_*A*_, *R, S*_*post*_, *D*_*s*_ in a 2 × 2 × (2 × 2 + 1) simulation study design:

1. *M*_*A*_ : Model as determined by the signal to noise ratio in the true model generating the data. Two values are: Signal : Noise = 1 : 1, which is equivalent to the statistical condition *E*(*Y*) : *σ* = 1 : 1, and Signal : Noise = 1 : 4, which is equivalent to the statistical condition *E*(*Y*) : *σ* = 1 : 1 in figure 6.

**Figure 6.**
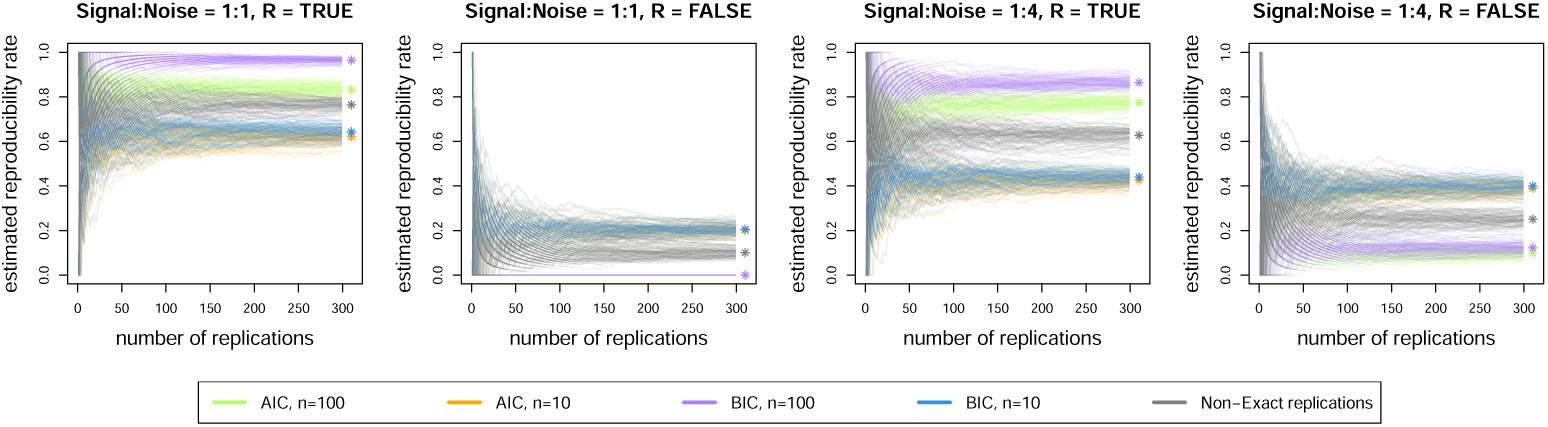
A simulation example to illustrate the convergence of reproducibility rates from exact and non-exact replication experiments to their true value. See text within the appendix for description of panels.
2. *R* : Result of the original experiment. Two values are: True and False.
3. *S*_*post*_ : Model selection method. Two values are: Akaike’s Information Criterion (AIC) and Bayesian Information Criterion (BIC).
4. *D*_*s*_ : Data structure. Two values are: Sample sizes *n* = 10 and *n* = 100.
5. Condition +1 : Uniformly randomly chosen non-exact replications at each step of the sequence from the set of all idealized experiments.

We performed 100 runs of a sequence of 1000 exact replication experiments for each of the sixteen experimental conditions, plus 100 runs of a sequence of 1000 non-exact replication experiments where (*M*_*A*_, *R, S*_*post*_, *D*_*s*_) is chosen uniformly randomly from sixteen conditions. Four of the experimental conditions (*M*_*A*_, *R* values) are shown in panels of figure 6: A: Signal : Noise = 1 : 1 and TRUE result; B: Signal : Noise = 1 : 1 and FALSE result; C: Signal : Noise = 1 : 4 and TRUE result; D: Signal : Noise = 1 : 4 and FALSE result. Five experimental conditions (*S*_*post*_, *D*_*s*_ values + non-exact replications) are shown in colored lines in each panel of figure 6. Green: AIC, *n* = 100; Orange: AIC, *n* = 10; Purple: BIC, *n* = 100; Blue: BIC, *n* = 10; Grey: non-exact replications. The result of interest is the reproducibility rate of the result of the original experiment, which is given by a star for each condition. The plots in figure 6 and the points to which they converge to illustrate how true reproducibility rate changes depending on the elements of *ξ* and the effect of divergence of *ξ′* from *ξ*. We emphasize that all parameters of the simulation example in figure 6 are chosen so that one can discern the effect of varying models, methods, and data structures.

We interpret the results as follows.

1. The reproducibility rates for false results and for true results sum to 1, which is a verification of simulation experiments.
2. By the true rates of reproducibility marked by stars, we observe that they depend on the true data generating mechanism, and the elements of the original experiment, *S*_*post*_ and *D*_*s*_. For example, as the noise increases, the true reproducibility rate gets smaller, and the variance of the estimated reproducibility rate increases. So for larger noise, replication results are expected to be highly variable. True reproducibility rates of true results also change with sample size and method.
3. Reproducibility rate increases with sample size for true results whereas it decreases for false results such that low sample size makes false results more reproducible in our simulations.
4. Even when the true reproducibility rate is high, we might see a lot of variation in observed reproducibility rate after a small number of replications even when they are exact replications. Fourth, non-exact replications yield highly variable observed reproducibility rates that do not converge to the true reproducibility rate of the original result.

This simulation experiment complements the one presented in the main text (figure 3) by providing a different illustration from our toy example. The context of linear regression models is readily relevant to many practicing scientists. Moreover, this simulation extends the results to new contexts by observing the outcome of interest under different levels of system noise and both true and false original results. Ultimately both simulations show considerable variability in true reproducibility rates as a function of the elements of and relationship between original and replication experiments.

## Appendix 9

### True results are not necessarily reproducible and perfectly reproducible results may not be true

Reproducibility is a function of the true unknown data generating model and the elements of *ξ*. Devezer et al. (2021) *provides some account. We give a brief overview with a proof by counterexample. Conditional on R* from *ξ*, we let *ξ*^(1)^, *ξ*^(2)^, … be exact replications of *ξ* and **I**_{*b*\*}_ be the indicator function that equals 1 if the first raven in the sample is black, and 0 otherwise. To prove the first part of the statement we choose the estimator

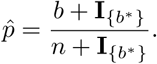

The estimator 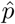 is valid on [0, 1] by: if *b* = *n*, then the first raven sampled must be black and 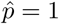, else if *b* = 0, then the first raven must be white and 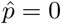 such that 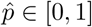. However, 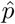 is unbiased for *p* only with probability (1 − *p*). The reason is that the probability of first raven is white raven is (1 − *p*) and if it is a white raven we get 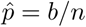 giving 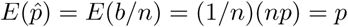. In contrast, 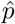 is biased for *p* with probability (1 − *p*). The reason is that the probability of first raven is black raven is *p* and if it is a black raven we get 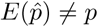. This does not only show that the true results are not always reproducible, but also shows that the reproducibility rate can be a function of the true parameter.

To prove the second part of the statement, choose the estimator 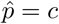, where *c* is a constant in [0, 1]. 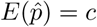. This expectation is only equal to *p* when *p* = *c*. However, the result using this 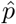 is reproducible with probability 1, thereby completing the proof.

## Footnotes

1 Some of the ideas developed in depth here appeared in preliminary form in (Baumgaertner et al., 2018).

2 We are not the first to take issue with the “replication crisis” framing. We invite the interested reader to visit Feest (2019)’s provocative and incisive assessment of why replication is overrated.

3 We assume that the order in which the data values appear has no bearing on the inferential goal. The cases in which the order contains information are important for a variety of subject matters, but it is well known that the statistical techniques that deal with them are too specialized to be treated in a general setup. An example is autoregressive models.

4 Compare this statement to definition 2 of an exact and non-exact replication experiment unconditional on an inferential objective.

5 Cooper and Guest (2014) and Guest and Martin (2021) make a similar point for computational reproducibility. They highlight the importance of making models available, and particularly clearly reporting model specifications and implementation assumptions so as to facilitate replication.

6 See Devezer et al. (2021) for examples of *ϕ* = 1 under uncertainty.

## References

Bak-Coleman, J., Mann, R. P., West, J., and Bergstrom, C. T. (2022). Replication does not measure scientific productivity.

Baribault, B., Donkin, C., Little, D. R., Trueblood, J. S., Oravecz, Z., van Ravenzwaaij, D., White, C. N., De Boeck, P., and Vandekerckhove, J. (2018). Metastudies for robust tests of theory. Proceedings of the National Academy of Sciences, 115(11):2607–2612.

Baumgaertner, B., Devezer, B., Buzbas, E. O., and Nardin, L. G. (2018). Openness and reproducibility: Insights from a model-centric approach.

Borgman, C. L. (2012). The conundrum of sharing research data. Journal of the American Society for Information Science and Technology, 63(6):1059–1078.

Botvinik-Nezer, R., Holzmeister, F., Camerer, C. F., Dreber, A., Huber, J., Johannesson, M., Kirchler, M., Iwanir, R., Mumford, J. A., Adcock, R. A., et al. (2020). Variability in the analysis of a single neuroimaging dataset by many teams. Nature, 582(7810):84–88.

Box, G. E. (1976). Science and statistics. Journal of the American Statistical Association, 71(356):791–799.

Bruns, S. B. and Ioannidis, J. P. A. (2016). P-curve and p-hacking in observational research. PLOS ONE, 11(2):1–13.

Chang, H. (2004). Inventing temperature: Measurement and scientific progress. Oxford University Press.

Collaboration, O. S. et al. (2015). Estimating the reproducibility of psychological science. Science, 349(6251):aac4716.#x2013;1–aac4716–8.

Collins, F. S. and Tabak, L. A. (2014). Policy: Nih plans to enhance reproducibility. Nature News, 505(7485):612–613.

Cooper, R. P. and Guest, O. (2014). Implementations are not specifications: Specification, replication and experimentation in computational cognitive modeling. Cognitive Systems Research, 27:42–49.

Dennis, B., Ponciano, J. M., Taper, M. L., and Lele, S. R. (2019). Errors in statistical inference under model misspecification: evidence, hypothesis testing, and aic. Frontiers in Ecology and Evolution, 7(372):1–28.

Devezer, B., Nardin, L. G., Baumgaertner, B., and Buzbas, E. O. (2019). Scientific discovery in a model-centric framework: Reproducibility, innovation, and epistemic diversity. PLOS ONE, 14(5):1–23.

Devezer, B., Navarro, D. J., Vandekerckhove, J., and Ozge Buzbas, E. (2021). The case for formal methodology in scientific reform. Royal Society Open Science, 8(3):200805.

Feest, U. (2019). Why replication is overrated. Philosophy of Science, 86(5):895–905.

Feyerabend, P. (1993). Against method. Verso.

Fletcher, S. C. (2021). How (not) to measure replication. European Journal for Philosophy of Science, 11(2):57.

Gelman, A. and Loken, E. (2013). The garden of forking paths: Why multiple comparisons can be a problem, even when there is no “fishing expedition” or “p-hacking” and the research hypothesis was posited ahead of time. Department of Statistics, Columbia University, 348.

Godden, D. R. and Baddeley, A. D. (1975). Context-dependent memory in two natural environments: On land and underwater. British Journal of psychology, 66(3):325–331.

Gruijters, S. L. (2022). Making inferential leaps: Manipulation checks and the road towards strong inference. Journal of Experimental Social Psychology, 98:104251.

Guest, O. and Martin, A. E. (2021). How computational modeling can force theory building in psychological science. Perspectives on Psychological Science, 16(4):789–802.

Hardwicke, T. E., Mathur, M. B., MacDonald, K., Nilsonne, G., Banks, G. C., Kidwell, M. C., Hofelich Mohr, A., Clayton, E., Yoon, E. J., Tessler, M. H., Lenne, R. L., Altman, S., Long, B., and Frank, M. C. (2018). Data availability, reusability, and analytic reproducibility: Evaluating the impact of a mandatory open data policy at the journal cognition. Royal Society Open Science, 5(8):1–18.

Hyman, I. (2021).

Iqbal, S. A., Wallach, J. D., Khoury, M. J., Schully, S. D., and Ioannidis, J. P. (2016). Reproducible research practices and transparency across the biomedical literature. PLoS biology, 14(1):1–13.

Janssen, M., Charalabidis, Y., and Zuiderwijk, A. (2012). Benefits, adoption barriers and myths of open data and open government. Information systems management, 29(4):258–268.

Kerr, N. L. (1998). Harking: Hypothesizing after the results are known. Personality and Social Psychology Review, 2(3):196–217.

Kuhn, T. S. (1970). The structure of scientific revolutions, volume 111. Chicago University of Chicago Press.

Lindley, D. V. (2000). The philosophy of statistics. Journal of the Royal Statistical Society: Series D (The Statistician), 49(3):293–337.

Loken, E. and Gelman, A. (2017). Measurement error and the replication crisis. Science, 355(6325):584–585.

Molloy, J. C. (2011). The open knowledge foundation: open data means better science. PLoS biology, 9(12):1–4.

Munafò, M. R., Nosek, B. A., Bishop, D. V. M., Button, K. S., Chambers, C. D., du Sert, N. P., Simonsohn, U., Wagenmakers, E.-J., Ware, J. J., and Ioannidis, J. P. A. (2017). A manifesto for reproducible science. Nature Human Behaviour, 1(0021):1–9.

Murre, J. M. (2021). The godden and baddeley (1975) experiment on context-dependent memory on land and underwater: a replication. Royal Society open science, 8(11):200724.

National Academies of Sciences, Engineering, and Medicine (2017). Fostering integrity in research. National Academies Press, Washington, D.C.

Nosek, B. A., Alter, G., Banks, G. C., Borsboom, D., Bowman, S. D., Breckler, S. J., Buck, S., Chambers, C. D., Chin, G., Christensen, G., et al. (2015). Promoting an open research culture. Science, 348(6242):1422–1425.

Nosek, B. A., Ebersole, C. R., DeHaven, A. C., and Mellor, D. T. (2018). The preregistration revolution. Proceedings of the National Academy of Sciences, 115(11):2600–2606.

Nosek, B. A., Hardwicke, T. E., Moshontz, H., Allard, A., Corker, K. S., Dreber, A., Fidler, F., Hilgard, J., Kline Struhl, M., Nuijten, M. B., et al. (2022). Replicability, robustness, and reproducibility in psychological science. Annual Review of Psychology, 73:719–748.

Rao, C. R. (1973). Linear statistical inference and its applications, volume 2. Wiley New York.

Serfling, R. J. (1980). Approximation theorems of mathematical statistics. John Wiley and Sons.

Shively, T. and Walker, S. (2013). On the equivalence between bayesian and classical hypothesis testing. arXiv preprint arXiv:1312.0302.

Silberzahn, R., Uhlmann, E. L., Martin, D. P., Anselmi, P., Aust, F., Awtrey, E., Bahník, Š., Bai, F., Bannard, C., Bonnier, E., Carlsson, R., Cheung, F., Christensen, G., Clay, R., Craig, M. A., Rosa, A. D., Dam, L., Evans, M. H., Cervantes, I. F., Fong, N., Gamez-Djokic, M., Glenz, A., Gordon-McKeon, S., Heaton, T. J., Hederos, K., Heene, M., Mohr, A. J. H., Högden, F., Hui, K., Johannesson, M., Kalodimos, J., Kaszubowski, E., Kennedy, D. M., Lei, R., Lindsay, T. A., Liverani, S., Madan, C. R., Molden, D., Molleman, E., Morey, R. D., Mulder, L. B., Nijstad, B. R., Pope, N. G., Pope, B., Prenoveau, J. M., Rink, F., Robusto, E., Roderique, H., Sandberg, A., Schlüter, E., Schönbrodt, F. D., Sherman, M. F., Sommer, S. A., Sotak, K., Spain, S., Spörlein, C., Stafford, T., Stefanutti, L., Tauber, S., Ullrich, J., Vianello, M., Wagenmakers, E.-J., Witkowiak, M., Yoon, S., and Nosek, B. A. (2018). Many analysts, one data set: Making transparent how variations in analytic choices affect results. Advances in Methods and Practices in Psychological Science, 1(3):337–356.

Stanley, D. J. and Spence, J. R. (2014). Expectations for replications: Are yours realistic? Perspectives on Psychological Science, 9(3):305–318.

Steegen, S., Tuerlinckx, F., Gelman, A., and Vanpaemel, W. (2016). Increasing transparency through a multiverse analysis. Perspectives on Psychological Science, 11(5):702–712.

Steinle, F. (1997). Entering new fields: Exploratory uses of experimentation. Philosophy of science, 64:S65–S74.

Stodden, V. (2011). Trust your science? open your data and code. Amstat News, July:21–22.

van’t Veer, A. E. and Giner-Sorolla, R. (2016). Pre-registration in social psychology—a discussion and suggested template. Journal of Experimental Social Psychology, 67:2–12.

Wagenmakers, E.-J., Wetzels, R., Borsboom, D., van der Maas, H. L., and Kievit, R. A. (2012). An agenda for purely confirmatory research. Perspectives on Psychological Science, 7(6):632–638.

